# Seizure recruitment properties are dependent upon dynamotype: a modeling study

**DOI:** 10.64898/2026.02.04.703690

**Authors:** Diana M. Karosas, Marisa Saggio, William C. Stacey

**Affiliations:** University of Michigan; Aix-Marseille Universite

## Abstract

Seizure propagation – how epileptogenic brain regions recruit less excitable regions – is poorly understood. Previous studies have used dynamical modeling to study seizure propagation and to create patient-specific whole-brain models of seizure spread. However, these studies focused on seizures of a single dynamotype (onset and offset bifurcation pair). Here, we implement a novel coupling method to investigate seizure propagation in a diverse array of dynamotypes. We utilize the Multiclass Epileptor, a recently proposed model that captures a wide range of seizure dynamotypes in a cortical mass (“node”). We consider two nodes: the seizure onset zone (node 1), which bursts autonomously, and the potential propagation zone (node 2), which is not independently epileptogenic but can be recruited by node 1. We examine the impact of intrinsic and coupling factors on the likelihood and speed of recruitment, with particular attention to the onset bifurcation of node 1. We also measure the range of onset behaviors observed in node 2 with respect to the onset behavior of node 1. The model predicted that seizures that display baseline shifts at onset are less likely to spread, and spread more slowly, compared to seizures that do not exhibit baseline shifts at onset. Seizures that present with amplitude scaling at onset were unlikely to propagate. Further, the model predicted the potential for unusual combinations of onset dynamics, such as a baseline shift in node 2 but not node 1. We confirmed the possibility for several of these unusual recruitment behaviors in humans using intracranial electroencephalography data. The results of the study provide a theoretical framework for seizure propagation, establishing a basis for innovations in characterization of patients’ seizure networks and identification of the seizure onset zone.

**Author Summary:** In this work, we examined how a seizure spreads from one part of the brain to another using a computational model. We modeled two brain regions using the Multiclass Epileptor, which reproduces a range of brain activity patterns associated with seizures. In the model, the first brain node was able to recruit the second brain node into a seizure. The model predicted that the likelihood and speed of seizure spread differ depending on the pattern of brain activity observed at the start of the seizure. We also found that the pattern of brain activity at seizure onset is not necessarily the same pattern seen when the seizure spreads. We confirmed this possibility for mismatched patterns in recordings from human brain. The findings of the study improve our understanding of seizure spread, which lays the groundwork for development of tools to quantify seizure spread and may inform future work in patient-specific brain modeling.

## 1. Introduction

Epilepsy, defined as the propensity for multiple seizures [1], affects over 50 million people and carries a global burden of disease of 13.9 million disability-adjusted life-years [2]. Antiseizure medications are the first-line treatment for epilepsy, but are insufficient to achieve seizure freedom in at least one-third of patients [3,4]. Surgical resection is a treatment option for drug-resistant focal epilepsy; however, between 20% and 60% of individuals continue to experience seizures following resective surgery [3,5]. Resection aims to remove the region of brain responsible for initiating the seizure, known as the epileptogenic zone [6]. However, seizures typically propagate beyond that region once the seizure begins. A contributing factor to ineffective surgical outcomes is a lack of understanding of seizure propagation.

The current gold standard for delineation of the seizure onset and propagation zones is visual intracranial EEG analysis by a clinical expert [7,8]. That determination is primarily based upon identifying the first channels that begin a seizure, known as the Seizure Onset Zone [7]. Clinical reports also describe how the seizure spreads, but such descriptions of seizure propagation are not standardized. Propagation is primarily assessed via patterns of latency (i.e. how long until a channel joins the seizure) and synchronicity (i.e. if the propagated channel is firing in synch with the onset channels). Reliance on visual interpretation restricts the definitive identification of onset, early, and late spread zones, leading to a limited understanding of seizure propagation. Clinicians utilize empirical descriptions to decide when propagation is “early” or “late” and how to interpret such results. A more robust method of quantifying seizure propagation would aid in better characterization of the seizure network.

Better understanding of propagation requires an appropriate model system that is relevant across a range of seizures. Epilepsy emerges from a multitude of underlying conditions, and findings from one patient or etiology may not necessarily generalize to others [3,9]. Focusing on invariant properties common to seizures across etiologies, rather than the specific mechanisms of a particular type of epilepsy, may yield more widely applicable results. One invariant property is how the brain transitions between resting state and seizure [10–12], which can be described using bifurcation theory from nonlinear dynamics. Previous studies that used large-scale brain models to investigate seizure propagation and surgical targets equipped each brain region with identical local dynamics, differing only in excitability, which defines the distance to the (shared) onset bifurcation [13–21]. In such modeling frameworks, the only discriminant between seizure onset, propagation, and non-epileptic zones is their proximity to the onset bifurcation and how coupling with other nodes modifies this distance. One model used to define local dynamics is the Epileptor, which mimics one combination of onset/offset bifurcations found in human seizures [11].

Epileptors have been coupled together to study seizure recruitment in a toy model [20] and in full reconstructions of an epileptic brain [19,21]. That approach has been very successful in describing human seizures with The Virtual Brain, a model that recapitulates the connectivity of an individual patient using nodes to represent brain regions, each equipped with an Epileptor model [21].

However, seizure onset and offset dynamics are heterogeneous among patients and can change from seizure to seizure in a single patient, calling for further investigation of the impact of local dynamics on seizure propagation. Previous studies on coupled nodes with different onset/offset bifurcations pairs are rare and focus on synchronization rather than recruitment [22]. The Multiclass Epileptor is capable of reproducing a wide range of local dynamics found in the Taxonomy of Seizure Dynamotypes, which classifies seizures based on their onset and offset bifurcations [11,12,23–25]. That work described four onset and four offset bifurcations, making 16 dynamotypes that encompass a wide range of dynamical behaviors. Heterogeneities compatible with different dynamotypes were assessed using seizures recorded via intracranial EEG in 120 patients with focal epilepsy [24], and this framework has since been used to quantify epileptogenesis in rats [26] and to predict responses to ictal-aborting stimulation [27].

The goal of this work is to provide the first exploration of recruitment behaviors when different brain regions have potentially different dynamical behaviors. The strategy is to describe behaviors across multiple local dynamics in the active seizure focus and the recruited brain region, with specific attention to conditions in which seizures are more or less likely to propagate. We implement a novel method to couple two simulated brain regions using the Multiclass Epileptor to enrich our understanding of recruitment of non-epileptogenic brain regions during a seizure. We first analyze the coupled model theoretically, then simulate recruitment for a variety of coupling strengths, onset zone dynamics, and propagation zone dynamics. These simulations indicate that seizure recruitment properties depend on (1) coupling strength; (2) the dynamics of the seizure onset zone; and (3) the resting state dynamics of the propagation zone. The majority of results show that the propagation zone is recruited to have similar dynamics to the onset zone. However, the model also predicts certain conditions in which the two have different dynamics and provides a mathematical explanation for this behavior. We provide examples of several of the different predicted recruitment behaviors with intracranial EEG recordings in patients with epilepsy. The results of the study provide a dynamical framework for seizure propagation, deepening our understanding of recruitment of non-epileptogenic brain regions during a seizure. Factors that influence recruitment likelihood and speed are identified, providing a basis for future innovations in the characterization of patients’ seizure networks and identification of the seizure onset zone.

## 2. Methods

### 2.1 The Multiclass Epileptor

A generic dynamical engine that reproduces a wide range of dynamotypes was previously created by exploiting the canonical structure around the degenerate Bogdanov-Takens singularity [12]. In the context of epilepsy, we refer to this DBT Burster as the Multiclass Epileptor to highlight that it is a more comprehensive version of the Epileptor, which describes one seizure dynamotype [11]. A full description of the DBT Burster has previously been published [12]; here, we provide a brief summary of the model to lay a foundation of understanding for our novel coupling method.

From a dynamical perspective, epilepsy is characterized by at least two rhythms: a fast rhythm governs neural activity during a seizure, while a slower rhythm or process governs the transitions between seizure and resting states. Bursting thus characterized by periods of quiescence and activity is termed ‘fast-slow bursting’, and can be described by Eqns. 1-2 [12,23]. Eqn. 1 describes the fast subsystem given by state vector 𝒙 = 𝒙(𝑡) ∈ ℝ^*n*^, and Eqn. 2 describes the slow subsystem given by state vector 𝒛 = 𝒛(𝑡) ∈ ℝ^*m*^. Timescale separation between the subsystems is ensured by setting 𝑘 ≪ 1.

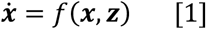

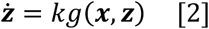

Reported in detail elsewhere [12,25,28], a minimally complex realization of the fast subsystem, comprised of two state variables such that 𝒙 = (𝑥, 𝑦), is given by Eqns. 3-4.

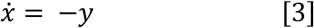

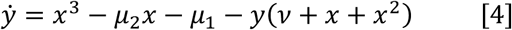

The state variable 𝑥 is the output of the system, and tracks voltage content during a seizure, the equivalent of an EEG signal. Within the state space of the model, resting states are represented by stable fixed points, and ictal states are represented by stable limit cycles. The existence of stable fixed points and limit cycles and therefore the behavior of the fast subsystem vary depending on the values of its parameters (𝜇_1_, 𝜇_2_, 𝜈). Although the model is phenomenological, these parameters have similar dynamical role to the influence of physiological factors such as ion concentrations, oxygen consumption, synaptic activity, etc. [11,24].

As in previous work [12,27], we constrain the parameters of the fast subsystem to the surface of a sphere centered at the origin with radius 0.4 (Fig.1). This reduces the dimensionality of the parameter space to two (the angular coordinates theta and phi). According to the location of the fast subsystem in parameter space, the system may be ‘resting’ (only a stable fixed point(s) exists), ‘ictal only’ (only a stable limit cycle exists), or ‘bistable’ with potential for both resting and ictal behavior (both a stable fixed point and a stable limit cycle exist). In the resting region, we distinguish ‘rest’ from ‘active rest’ based on the location of the fixed point in state space. The difference in the location of the fixed points is due to the geometry of the equilibrium manifold, as explained previously [12,24]. Active rest occurs in the top-right quadrant of the parameter space sphere; all other resting regions are rest. Two rest-ictal bistability regions, with distinct state space topologies, exist on the surface of the parameter space sphere. In the ‘limit cycle small’ bistability region (LCs), the stable fixed point is separated from the limit cycle. Due to this separation, transitions from stable fixed point to limit cycle and vice versa are associated with baseline shifts in the 𝑥 variable. In the ‘limit cycle big’ bistability region (LCb), the stable fixed point is within the limit cycle. Thus, transitions between the stable fixed point and limit cycle in LCb are not associated with baseline shifts.

**Fig 1.**
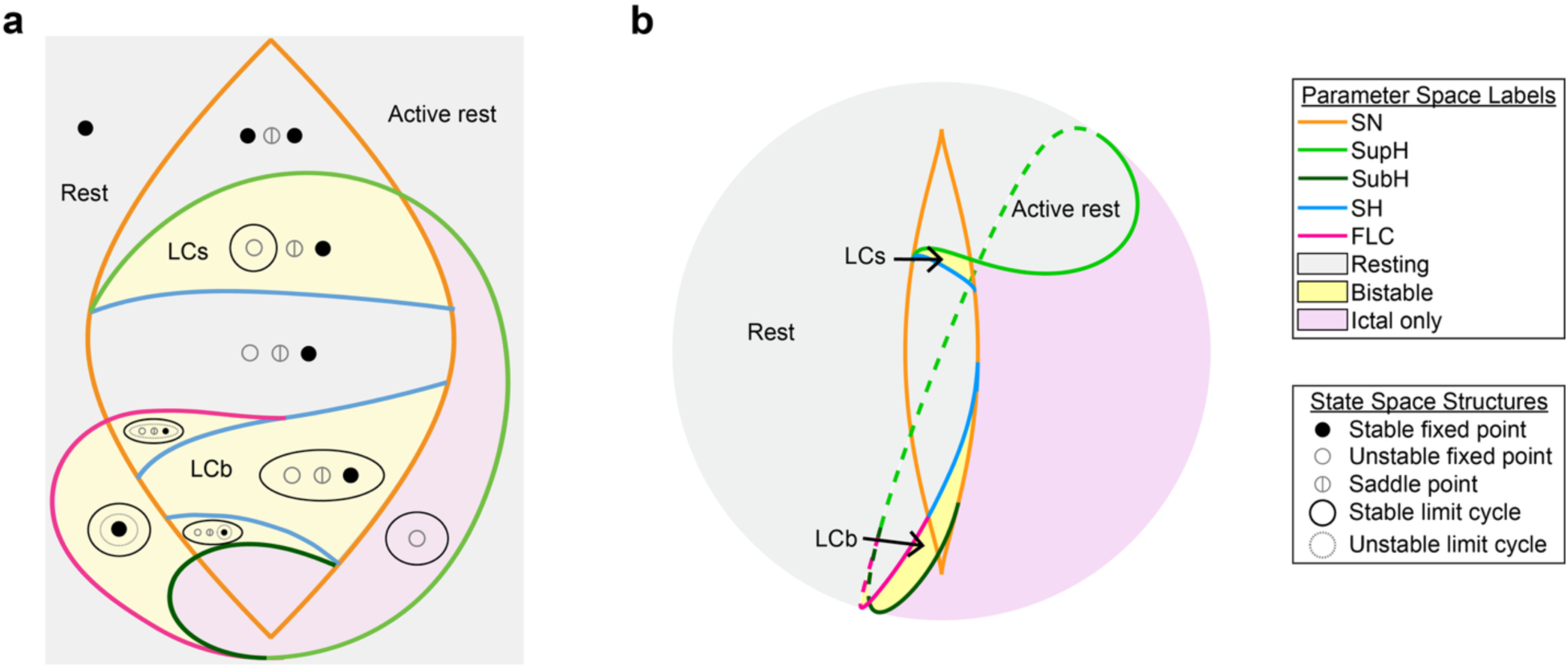
**(a)** 2D illustration and **(b)** 2D projection (lambert equal area) of parameter space. Bifurcation curves (colored lines) separate qualitatively distinct behavioral regions (shaded). Dashed lines indicate the back of the sphere in (b). State space diagrams are drawn in each region in (a). ‘Active rest’ includes the ‘Resting’ region to the right of the SN bifurcation in the top half of the sphere; all other ‘Resting’ regions are ‘Rest’. Two bistability regions exist: limit cycle small (‘LCs’), where the stable fixed point is outside the limit cycle; and limit cycle big (‘LCb’), where the stable fixed point is inside the limit cycle. SN = saddle node, SupH = supercritical Hopf, SubH = subcritical Hopf, SH = saddle homoclinic, FLC = fold limit cycle.

There are two ways to create a burst or seizure in the model [10,12], which link to the two major dynamical mechanisms proposed for seizure onset at the level of a single node [18,29,30]. Within the bistability regions, adding noise or an appropriate stimulus can cause the system to move between stable fixed point and stable limit cycle. The parameters of the fast subsystem need not change for such noise-induced transitions, although proximity to an onset bifurcation lowers the threshold for the transition.

Alternatively, the transitions between resting and active states can be precipitated by slow changes in the fast subsystem’s parameters that alter the existence of stable fixed points and limit cycles; these types of transitions are known as bifurcations. On the surface of the parameter space sphere, bifurcations are represented by curves that divide regions of qualitatively distinct behavior repertoires (Fig.1). A seizure can be created when the fast subsystem traverses a path that crosses onset and offset bifurcations in parameter space. The slow subsystem of Eqn. 2 moves the fast subsystem along this bursting path.

Multiple realizations of the slow subsystem are possible [12,23,25]. In particular, hysteresis-loop implementations exploit bistability in the fast subsystem and feedback among fast and slow variables to produce autonomous bursting. Slow-wave implementations lack this feedback, may occur without bistability, and require slow variables to change independently to force the fast ones into bursting. For the present work, we chose hysteresis-loop paths for the node representing the active seizure focus (node 1) because it allows this node to start and end a seizure autonomously. A single implementation was chosen for comparison among bursting paths.

Eqn. 5 describes the speed and direction of movement of node 1 along its bursting path. In Eqn. 5, 𝐷 represents the Euclidean distance (in (𝑥, 𝑦) state space) from rest and 𝑑^∗^ represents the excitability of the system. A larger excitability allows the fast system to move further from the resting state before the direction of motion in parameter space changes from toward onset bifurcation to toward offset bifurcation. Increasing 𝑑^∗^ alters the duty cycle of the slow subsystem such that the fast subsystem spends a greater proportion of time in the active (vs quiescent) state and near the onset bifurcation. The speed at which the fast subsystem moves along the bursting path is determined by 𝑘_1_. To maintain timescale separation as in Eqn. 2, 𝑘_1_ << 1. Eqn. 5 prescribes that the system moves toward onset bifurcation when it is at rest and moves toward offset bifurcation when it is bursting.

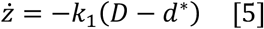

A necessary condition for Eqn. 5 is that the bursting path exists in a bistable region of parameter space. For simplicity and without loss of generality, we choose the bursting paths to be arcs of great circles that connect onset and offset bifurcations. Details of the implementation method can be found in [12]. The specific onset and offset bifurcations determine the dynamotype of the burst or seizure. There are eight dynamotypes that can be implemented on this sphere using the hysteresis mechanism. We separated these eight dynamotypes into four groups by onset bifurcation in this work: saddle node with baseline shift (SN(+DC)), supercritical Hopf (supH), saddle node without baseline shift (SN(-DC)), and subcritical Hopf (subH). The invariant characteristics of these onset bifurcations are indicated in Table 1 [12,24,25].

**Table 1:** Onset bifurcation characteristics.

| Bifurcation | Amplitude scaling at bifurcation point | Frequency at bifurcation point | Baseline shift in x at bifurcation point |
| --- | --- | --- | --- |
| Saddle node with baseline shift (SN(+DC)) | no | nonzero | yes |
| Supercritical Hopf (supH) | yes | nonzero | no |
| Saddle node without baseline shift (SN(-DC)) | no | nonzero | no |
| Subcritical Hopf (subH) | no | nonzero | no |

### 2.2 Connecting seizure onset and propagation zones

The Multiclass Epileptor describes the behavior of a cortical mass or “node”. Although the model describes seizure features that are present at different scales [11], it may be helpful to think of a node as the mass under a single SEEG electrode contact. To study seizure propagation, we coupled two nodes of the model. Clinical observations guided the development of our coupling method. When a seizure propagates, the propagation zone often exhibits the same onset pattern as and synchronizes with the seizure onset zone. However, the propagation zone does not always adopt the same dynamics as the seizure onset zone, and seizures often propagate to brain regions that are not independently epileptic. We could think of two parameter space possibilities to capture these tendencies. First, the highly excitable onset zone and less excitable propagation zone could inhabit similar regions of parameter space, and the onset zone could then influence the excitability of the propagation zone. Second, they could exist in different regions, and the onset zone could move the propagation zone into its region of parameter space during a seizure. The former explanation would require every node in the brain to inhabit the same, small, bistable region of parameter space. In such a case, all brain nodes would have similar underlying dynamics. Only the latter explanation provides a mechanism for recruitment of the propagation zone from multiple regions of parameter space, including resting regions. We chose the latter case. Thus, in our coupling method, we assume that a recruiting brain region changes the dynamics of the propagation zone to be similar to its own, but that the propagation zone is not required to initially be in the same region of parameter space.

Node 1 represented the seizure onset zone and was modeled as an unmodified hysteresis-loop burster using Eqn. 3-5. Since we were interested in the initial recruitment of brain regions into a seizure (and not complex feedback effects after a seizure starts such as suppression or patterns of seizure progression and termination), we implemented one-directional coupling. In other words, node 2 was affected by node 1 but node 1 was not influenced by node 2. While this is an oversimplification, it captures the underlying pathology of seizure onset activity, defined by when uncontrolled bursting overrides normal brain activity. Node 2 represented the potential propagation zone and was not independently epileptogenic. The fast subsystem of node 2 was unchanged from Eqn. 3-4. The slow subsystem was modified to allow the fast subsystem of node 1 to drive recruitment of node 2. Fast-to-slow coupling has been proposed as a mechanism for seizure propagation since it allows fast activity in the seizure focus to influence the slow variables in another brain region [19–21]. Since node 2 does not burst autonomously and only moves toward seizure onset according to input from its coupling with node 1, node 2’s bursting mechanism is effectively the slow-wave type.

In the propagation model, node 2 is initially placed at rest at a stationary point in parameter space, i.e. 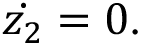 Upon the onset of the burst in node 1, node 2 begins to move toward node 1 in parameter space along an arc connecting the nodes, with speed proportional to the square of the difference between the current state of node 1 and its resting state (in 𝑥). Since the x variable mimics a voltage recording, this form of coupling captures the influence of the change in voltage in the seizure onset zone (node 1) on the activity in the potential propagation zone (node 2). Following the offset of the burst in node 1, node 2 moves back to its initial location in parameter space at a constant speed (𝑘_2_). The parameterization of node 2’s path in parameter space, 𝝁(𝑧), is given by Eqns. 6-8. In these equations, **A** is the location of node 2. During the coupling period, **B** is the location of node 1; after coupling ends, **B** is the initial location of node 2. **A** and **B** are updated at every timestep. R is the radius of the sphere (0.4).

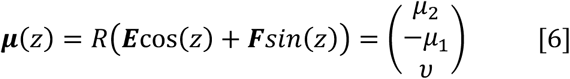

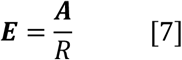

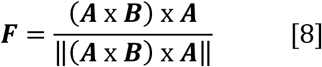

The speed of node 2’s movement along this path, 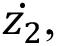 is defined according to Eqn. 9. In Eqn. 9, 𝑡_on,1_ is node 1 onset time, 𝑡*_off_*_,1_ is node 1 offset time, 𝑡_rest,2_ is the time when node 2 reaches its initial location in parameter space following the burst in node 1, and 𝐺 is a coupling strength constant.

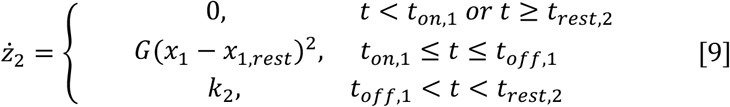

A representative example of recruitment showing the parameter space movement of the nodes and corresponding model outputs is provided in Fig.2 and Supplementary Video 1.

**Fig 2.**
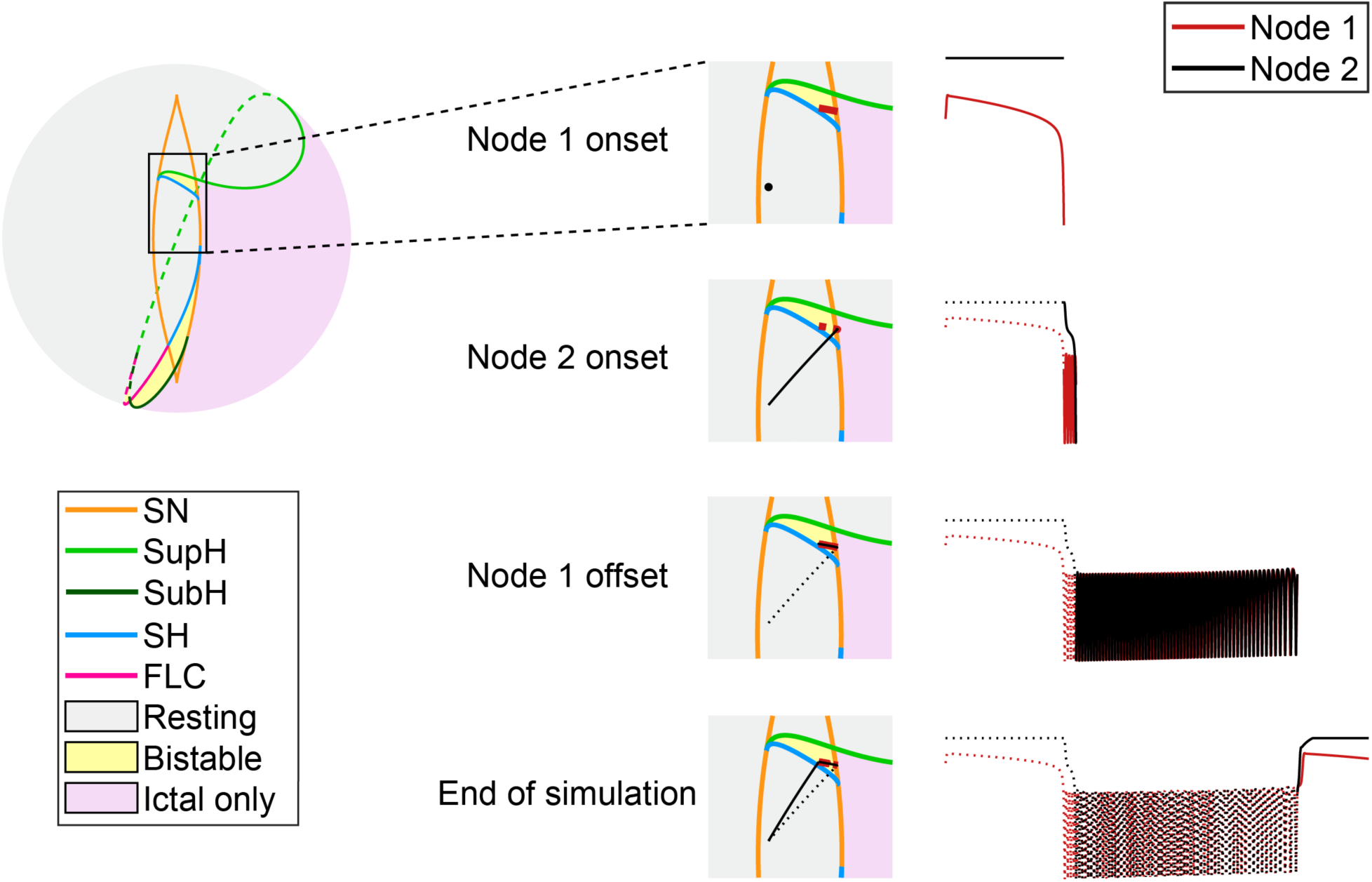
Example of recruitment of node 2 by node 1, showing parameter space movement (left) and model output (right) for each node. Node 1 begins to burst in the LCs region, while node 2 is far away in a resting region. During the burst, node 1 pulls node 2 towards itself, and node 2 starts to burst when it crosses the onset bifurcation curve. Results are shown at four time points in the simulation (top to bottom). In each timeseries, solid lines indicate activity since the last time point. SN = saddle node, SupH = supercritical Hopf, SubH = subcritical Hopf, SH = saddle homoclinic, FLC = fold limit cycle.

**Fig 3.**
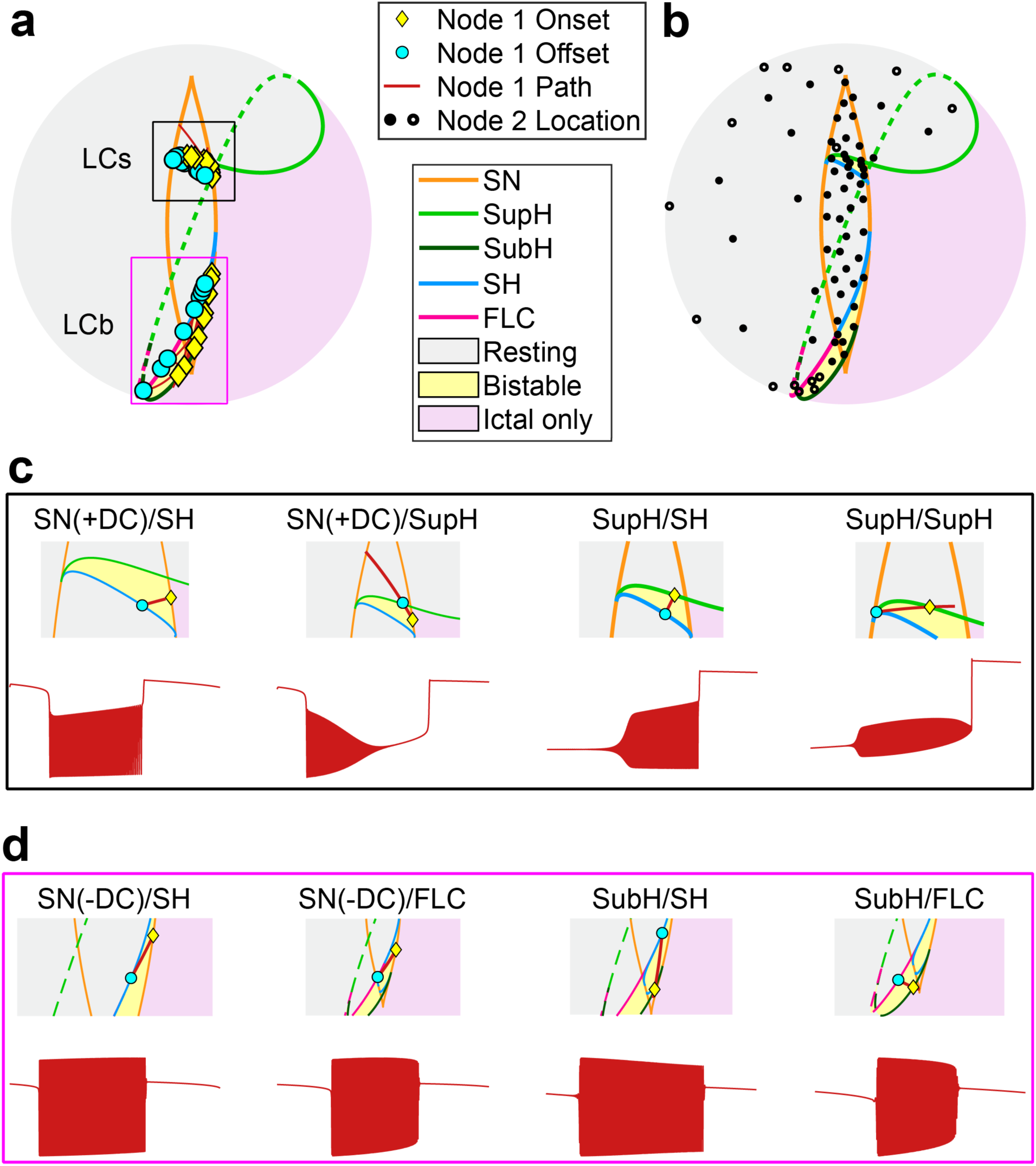
**(a)** Hysteresis-loop onset and offset locations and paths used to model node 1. **(b)** Parameter space locations used to model node 2. Open circles and dashed lines are used to indicate a location on the back of the sphere. **(c)** and **(d)** Zoomed-in views of a representative bursting path from (a) and timeseries, by limit cycle region and dynamotype class. LCs = limit cycle small, LCb = limit cycle big, SN(+DC)= saddle node with baseline shift, SN(-DC) = saddle node without baseline shift, SupH = supercritical Hopf, SubH = subcritical Hopf, SH = saddle homoclinic, FLC = fold limit cycle.

### 2.3 Simulation parameters

We simulated a wide range of node 1 classes, node 2 initial locations, coupling strength (𝐺) values, and node 1 excitability (𝑑^∗^) values to investigate the effects of these factors on recruitment properties. For each of the eight dynamotype classes that can be implemented via hysteresis-loop bursting, three paths were chosen arbitrarily. The speed of movement along each bursting path (𝑘_1_) was set slow enough that the characteristic properties of the dynamotype were evident in the timeseries. Initial conditions were adjusted such that the nodes were initially at rest. 𝑨, 𝑩, 𝑘_1_, and the initial conditions for each node 1 path are listed in Supplementary Table 1. The variability of ictal inter-spike intervals across bursting paths was confirmed to be within the expected physiological range (within an order of magnitude; Fig.S1). For each dynamotype, onset and offset bifurcations, parameter space bursting paths, and a representative timeseries are displayed in Fig.3a, c-d. Note that while we included all dynamotypes possible on the map in our analyses, offset bifurcation is less relevant to recruitment. Thus, our results are presented with respect to onset bifurcation only.

As illustrated in Fig.3b, 71 initial node 2 locations were simulated. Regions near bifurcations were sampled more heavily, while there was more sparse sampling in resting regions far from bifurcations. When initially located in a resting region, the system is in the quiescent state until a bifurcation is crossed, so there are only trivial differences in recruitment properties between two nearby points that are both far from bifurcations. Since node 2 was initially placed in the quiescent state (on a stable fixed point) to model a potential propagation zone, the ictal-only region of the sphere was avoided.

Seventeen values of coupling strength (𝐺) were tested between 0.001 and 0.3. Three node 1 excitability (𝑑^∗^) values were tested between 0.3 and 0.5. Together with the 24 bursting paths and 71 node 2 locations, there were a total of 86,904 simulations. For all simulations, the model was solved numerically using Euler’s forward method and a timestep of 0.01. Initial conditions were chosen such that the nodes were initially at rest. Timeseries were generated from the state variable 𝑥. All simulations lasted for a single burst in node 1. MATLAB R2024b was used for all simulations.

### 2.4 Recruitment analysis

#### 2.4.1 Identification of recruitment

For each simulation with at least 10 peaks (spikes), timeseries were high-pass filtered. Mean-square power was calculated for each node during the period with coupling between the nodes. If the node 2 power was at least 10% of the node 1 power, then node 2 was said to be recruited. This thresholding step was performed to discard node 2 responses that were not bursts. The threshold of 10% was chosen based on the observation that the proportion of simulations that resulted in recruitment was relatively stable for power thresholds between 0.1 and 0.8 (Fig.S2). Timeseries with power between 0.001 and 0.8 were visualized to confirm the choice of threshold was reasonable.

#### 2.4.2 Identification of burst onset and offset

In this work, we defined burst/seizure onset as the onset of sustained oscillations. In general, burst onset and offset in node 1 were marked by inspecting the sign of the derivative of the slow variable, ż. In the hysteresis-loop implementation, ż changes from positive to negative when node 1 departs from the stable fixed point representing rest; the opposite is true at burst offset (i.e. when node 1 returns to rest). For node 1 dynamotypes with a supH onset bifurcation, hysteresis requires a SN(+DC) bifurcation to move the system into the correct position in state space prior to the onset supH bifurcation. The sign of ż changes after the SN(+DC) bifurcation, but bursting does not start until the supH bifurcation. To isolate the recruitment properties of the supH bifurcation, we adjusted the initial conditions such that the first supH-onset burst could occur without a preceding SN bifurcation.

Specifically, we placed the node on the fixed point corresponding to active rest at the start of the simulation. We then defined burst onset as the first negative peak in the timeseries. Thus, node 1 did not influence node 2 through coupling (Eqn. 9) until the supH bifurcation. It should be noted that it is possible to observe periodic bursts in the Multiclass Epileptor that display a supH onset bifurcation without a preceding SN(+DC) bifurcation using paths other than the hysteresis-loop.

In node 2, burst onset was defined as the first negative peak in the timeseries after the onset of coupling. Identification of offset was not required for the analysis. For identification of peaks, MATLAB’s *findpeaks()* was used with a minimum peak prominence of 0.005. Peak prominence is a measure of how much a peak stands out from the surrounding baseline; an arbitrarily low value was set to ensure peaks due to numerical noise were ignored. Peaks that were not prominent and temporally separated from the rest of the bursting pattern were also ignored to remove transients that could occur at the onset of coupling. Peaks were considered ‘not prominent’ if they were below the mean prominence for the simulation. The mean prominence was calculated after discarding the bottom and top 10% to reduce the influence of outliers. To confirm the accuracy of onset times, a third of the data were visualized. Onset was confirmed to be marked accurately in over 99.9% of cases.

#### 2.4.3 Recruitment properties

Three recruitment properties were measured: the proportion of simulations that resulted in recruitment, recruitment delay, and recruitment energy. Recruitment delay was defined as the time between node 1 onset and node 2 onset. Recruitment energy was defined as the area under the recruitment term 𝐺(𝑥_1_ – 𝑥_*1,rest*_)^2^, measured from node 1 onset until node 2 onset. Recruitment energy captures the amplitude and duration of bursting in node 1 required to recruit node 2. For delay and energy calculations, time was divided by the duration of the burst in node 1 for comparison across node 1 dynamotypes and excitability values.

Results were presented with respect to node 1 onset bifurcation. There were four onset bifurcation groups: SN (+DC), supH, SN(-DC), and subH. To evaluate the effect of propagation zone dynamics on recruitment properties, analyses were repeated with respect to node 2 locations. Except where otherwise specified, the mean and variance of each measure were computed for each node 2 location, then normalized by subtracting the minimum and dividing by the range for visualization of trends.

### 2.5 Clinical data

Data were collected from a previously-deidentified database at the University of Michigan of long-term intracranial video-EEG monitoring in patients with epilepsy. The sampling rate was 4096 Hz. All patients consented to have their deidentified EEG data saved for research, and the protocol was approved by the local IRB. Seizure onset and propagation zones were identified by reading the official EEG report written by the treating clinicians.

## 3. Results

### 3.1 Theoretical Analysis

The Multiclass Epileptor is a dynamically rich, well-understood model of bursting that enables theoretical analysis. Here, using the parameter and state space topologies of the Multiclass Epileptor as a guide, we identify possible mechanisms of recruitment of node 2. We then predict dynamical conditions that encourage recruitment, and the behavior of node 2 when it is recruited. Note that while this analysis is provided in the context of coupled nodes of the Multiclass Epileptor, our approach and the recruitment mechanisms identified are applicable to other dynamical models of bursting.

#### 3.1.1 Possible mechanisms of recruitment

In this work, all node 1 bursting paths are located within bistability regions to facilitate autonomous bursting. Node 1 can enter bursting via SN(+DC) or supH bifurcation in LCs and via SN(-DC) or subH bifurcation in LCb (Fig. 4a, left). Note that only parts of the SN and supH bifurcations can induce bursting.

**Fig 4.**
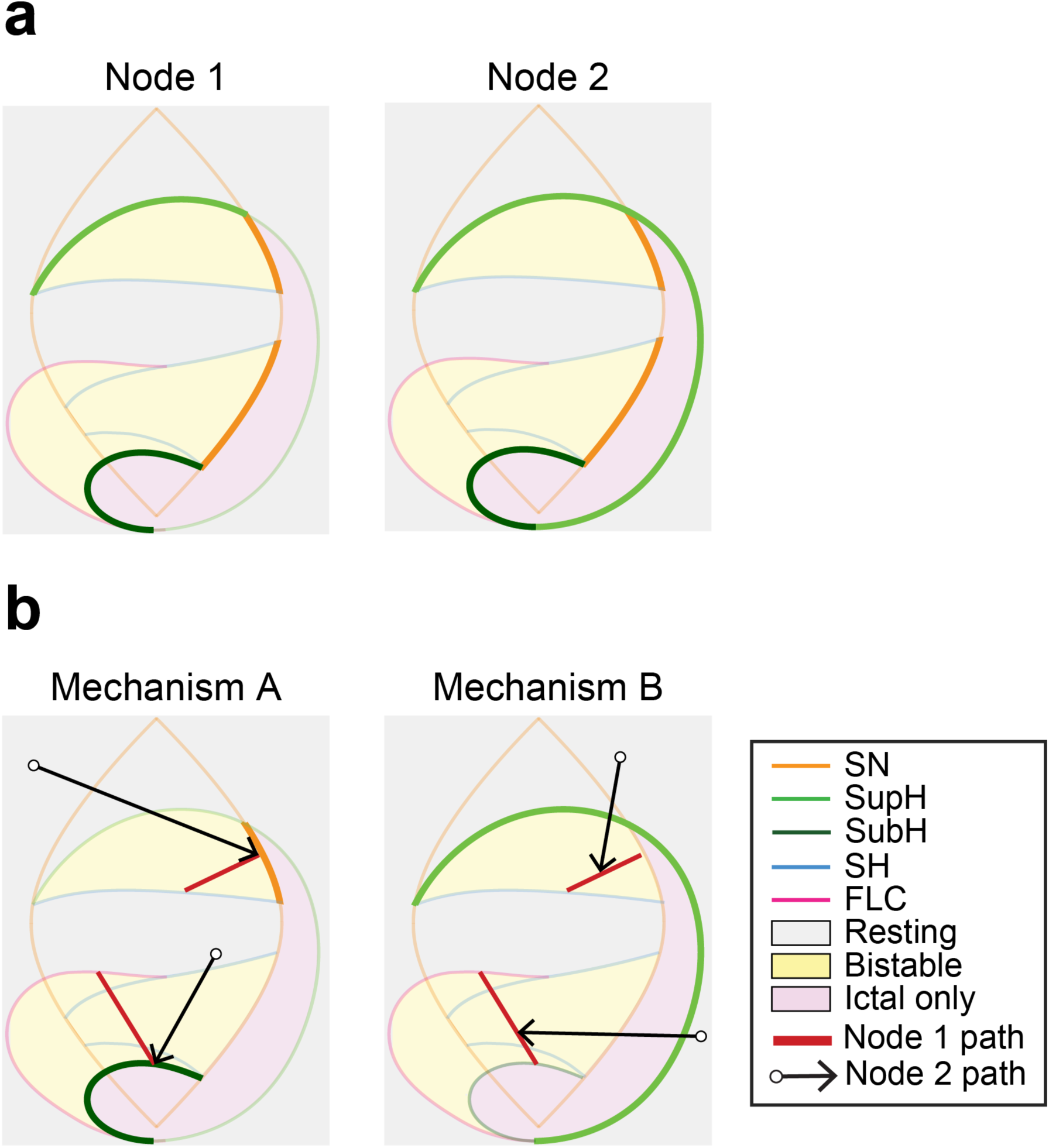
**(a)** Illustration of potential onset bifurcations for node 1 (left) and node 2 (right). Bifurcations that can serve as onset bifurcations are highlighted. **(b)** Illustration of two mechanisms of recruitment. Onset bifurcations for node 2 are highlighted. In mechanism A (left), node 2 (black) reaches node 1 (red) before node 1 leaves its onset bifurcation. In mechanism B (right), node 2 is recruited by crossing a bifurcation before reaching node 1. SN = saddle node, SupH = supercritical Hopf, SubH = subcritical Hopf, SH = saddle homoclinic, FLC = fold limit cycle.

Node 2 may initially be located anywhere in parameter space with the exception of the ictal-only region. Upon the onset of coupling, node 2 travels toward either the LCs or LCb bistability region, depending on the class of node 1. Two mechanisms exist by which node 2 can enter bursting (Fig. 4b). In mechanism A, node 2 does not cross onset bifurcations before reaching node 1 and reaches node 1 before node 1 has left its onset bifurcation. Node 2 can be recruited if node 1’s onset bifurcation is SN or subH since these bifurcations are embedded in the ictal-only region. Recruitment through mechanism A must occur early in the burst in node 1. Except in cases where the initial parameters of node 2 are very close to the onset bifurcation of node 1, mechanism A requires node 2 to temporarily break timescale separation, meaning the slow variable z is effectively moving at the same time scale as x. Mechanism A is more likely to occur when the coupling factor (𝐺) is high, which increases the speed of node 2’s movement toward node 1, and when node 1 excitability (𝑑^∗^) is high, which increases the relative amount of time node 1 spends near onset bifurcation after having crossed it. Examples of recruitment through mechanism A are given in Fig. 4b, left.

In mechanism B, node 2 is recruited by crossing an onset bifurcation along the path to, but before reaching, node 1. We can think of two scenarios for mechanism B. First, node 2 may pass through the ictal-only region along the path to node 1. The portion of the supH bifurcation that borders the ictal-only region can thus act as an onset bifurcation for node 2. Second, for some initial node 2 parameters, crossing the supH bifurcation to enter the LCs region promotes recruitment (see 3.1.2 and Fig. 5). In contrast to mechanism A, mechanism B can facilitate recruitment at any point in the burst in node 1.

**Fig. 5.**
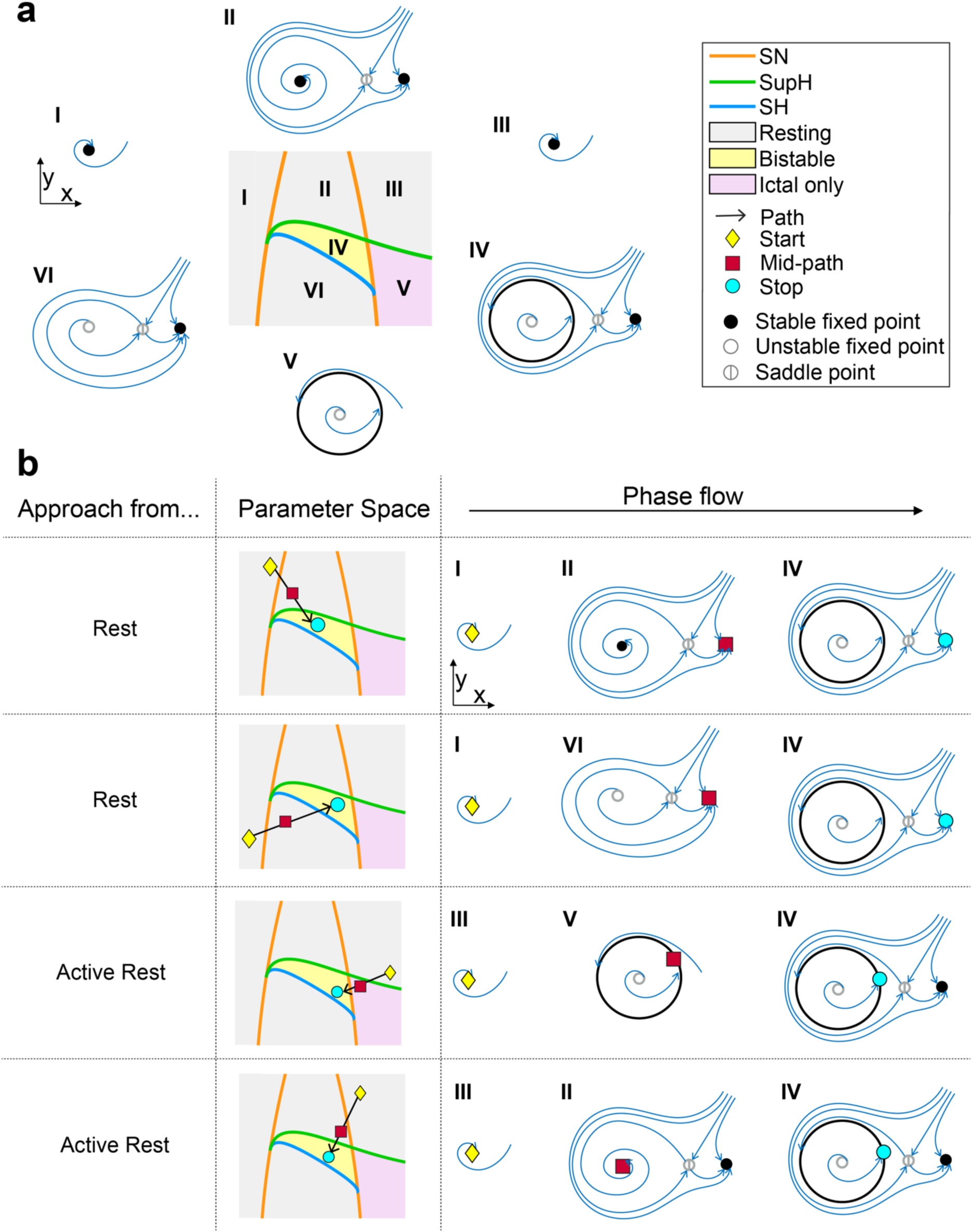
**(a)** Flow diagrams in each region surrounding LCs, and in LCs. **(b)** Phase flow diagrams demonstrating changes in state space during recruitment to LCs from different directions. Blue lines are flow lines. During recruitment from rest (top two panels), the system begins on a stable fixed point that remains stable along the path to and in the LCs region. During recruitment from active rest (bottom two panels), the system begins on a stable fixed point that becomes unstable along the path to or upon entering the LCs region. LCs = limit cycle small, SN = saddle node, SupH = supercritical Hopf, SH = saddle homoclinic.

Given that mechanism B does not rely as heavily on node 2’s speed compared to mechanism A, recruitment through mechanism B should also be less sensitive to coupling strength (𝐺) and node 1 excitability (𝑑^∗^). Examples of recruitment through mechanism B are given in Fig. 4b, right. Considering both recruitment mechanisms, the onset bifurcations that can induce bursting in node 2 include all those that can induce bursting in node 1, plus the portion of the supH bifurcation bordering the ictal-only region (Fig. 4a, right).

#### 3.1.2 Theoretical predictions

Three predictions arise from the preceding analysis. First, nodes in the LCb bistability region will be more likely to recruit than nodes in the LCs bistability region. The onset bifurcation curves in LCb bistability region are fully embedded in the ictal-only region, whereas only the SN(+DC) bifurcation in the LCs bistability region borders the ictal-only region (Fig. 1, 4a). The consequences of this topology are twofold: (1) supH-onset node 1 dynamotypes, which occur in LCs, cannot recruit through mechanism A and (2) node 2 is more likely to encounter the ictal-only region when approaching the LCb bistability region (mechanism B). Thus, recruitment through either identified mechanism is more likely when node 1 is in the LCb bistability region.

Second, nodes in active rest are more likely to be recruited than nodes in rest. Fig. 5 illustrates state space arguments for higher recruitment proportions when node 1 is in LCs and node 2 is in active rest compared to rest. If node 2 is initially in rest (Fig 5b, top two panels), it may enter LCs through the region above the supH bifurcation (region II in Fig.5) or the region below the SH bifurcation (region VI in Fig.5). Along either route, the stable fixed point corresponding to rest does not disappear, so recruitment is not guaranteed. Note that recruitment could still occur through mechanism A. In contrast, if node 2 begins in active rest, it may enter the LCs region via the ictal-only region or via the region between the SN bifurcations (region II in Fig.5). In either case, the stable fixed point corresponding to active rest becomes unstable and a stable limit cycle appears (Fig. 5b, bottom two panels). The location of the unstable fixed point is within the basin of attraction of the limit cycle, enabling recruitment through mechanism B.

Fig. 6 illustrates state space arguments for higher recruitment proportions when node 1 is in LCb and node 2 is in active rest compared to rest. In the coupling method employed, node 2 travels toward node 1 in parameter space along the shortest arc connecting the nodes. If node 2 begins in active rest, the arc connecting node 1 and node 2 passes through the ictal-only region of the sphere (Fig. 6b, bottom panel), guaranteeing recruitment through mechanism B. When node 2 begins on the stable fixed point corresponding to rest, it may approach LCb without entering the ictal-only region (Fig. 6b, top two panels). Recruitment can then only occur through mechanism A.

**Fig. 6.**
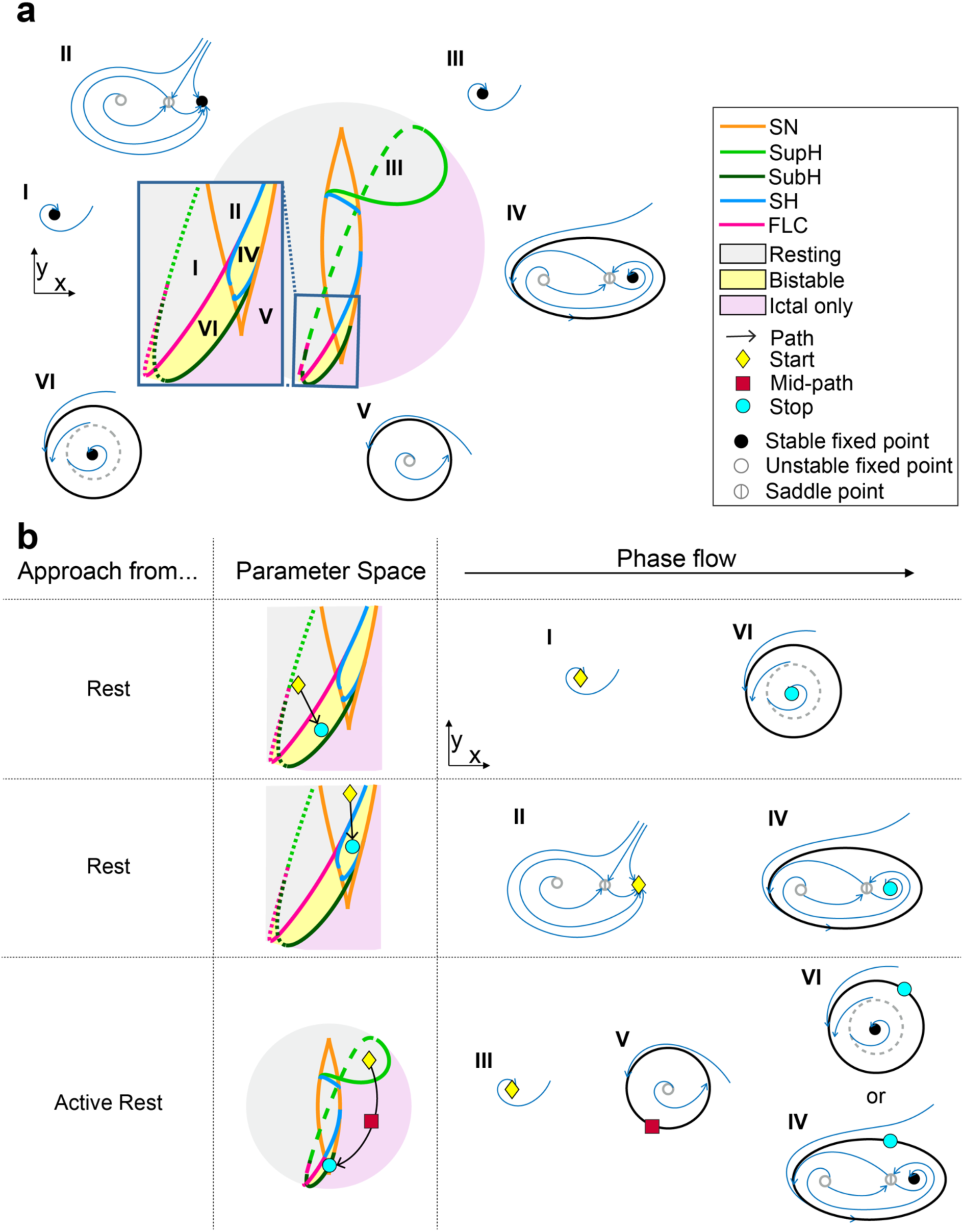
**(a)** Flow diagrams in each region surrounding LCb, and in LCb. **(b)** Phase flow diagrams demonstrating changes in state space during recruitment to LCb from different directions. Blue lines are flow lines. During recruitment from rest (top two panels), the system begins on a stable fixed point that remains stable along the path to and in the LCb region. During recruitment from active rest (bottom panel), the system begins on a stable fixed point that becomes unstable along the path to the LCb region. LCb = limit cycle big, SN = saddle node, SupH = supercritical Hopf, SubH = subcritical Hopf, SH = saddle homoclinic, FLC = fold limit cycle.

The third theoretical prediction is that the onset dynamics of node 2 may be the same as or different from those of node 1. Mechanism A ensures node 1 and node 2 encounter the same onset bifurcation, encouraging similar onset behavior. If node 1 exhibits a baseline shift, for example, node 2 will also exhibit a baseline shift. However, mechanism B allows for the dynamics of node 2 to diverge from those of node 1. When node 2 is recruited through mechanism B, it can exhibit supH or subH onset dynamics. Depending on node 1’s onset bifurcation, the dynamics of node 1 and node 2 may differ. Note that this prediction is only relevant with our choice of coupling mechanisms in which node 2 can initially be in a different region of parameter space than node 1 (see Section 2.2).

### 3.2 Dynamical factors that influence seizure recruitment *in silico*

Next, we simulated a wide range of dynamics in both the seizure onset zone and propagation zone [31]. We investigated recruitment properties via measurement of recruitment proportion, recruitment delay, and recruitment energy. We considered two dynamical factors that could influence recruitment properties: the onset dynamics of node 1 and the initial parameter space location of node 2. We also considered the effects of coupling strength (𝐺) and node 1 excitability (𝑑^∗^). Factors that were associated with higher recruitment proportions, lower recruitment delays, and/or lower recruitment energy were indicative of an increased tendency for recruitment. Finally, we identified the range of onset dynamics possible in node 2 for each node 1 onset type.

#### 3.2.1 Recruitment properties with respect to node 1 onset bifurcation

Node 1 dynamotypes were divided into four groups by onset bifurcation: SN(+DC), supH, SN(-DC), and subH. SN(+DC) onsets are characterized by baseline shifts. SupH onsets are characterized by increasing amplitude from zero. SN(-DC)-onset dynamotypes and subH-onset dynamotypes are both characterized by arbitrary amplitude without baseline shift [24]. Results with respect to node 1 onset bifurcation are summarized in Table 2.

**Table 2:** Summary of Recruitment Properties by Onset Bifurcation.

| Onset Bifurcation | Relative Recruitment Properties |  |  | Impact on Recruitment Likelihood |  |
| --- | --- | --- | --- | --- | --- |
| | Likelihood | Speed | Energy | Coupling Strength $G$ | Node 1 Excitability $d^*$ |
| SN (+DC) | Medium-Low | Slow | High | Medium | Medium |
| SupH | Low | Variable | Low | None | None |
| SN(-DC) | Medium-High | Fast | High | High | Medium |
| SubH | High | Fast | Medium | High | Low |

Consistent with theoretical predictions, nodes in the LCb bistability region were more likely to recruit than nodes in the LCs bistability region. SN(-DC) and subH-onset dynamotypes, located in LCb, showed elevated and faster recruitment (Fig.7a-b) compared to SN(+DC)-onset dynamotypes, located in LCs.

Recruitment proportion was particularly high for subH-onset dynamotypes. Recruitment energy was similar between SN-onset dynamotypes and lower for subH-onset dynamotypes, indicating that the disparities in recruitment proportion and delay were not due to increased separation from the resting state in subH-onset dynamotypes. Thus, the model predicts that seizures that do not start with baseline shifts are more likely to spread, and to spread faster, than seizures that display baseline shifts at onset.

Further, supH-onset dynamotypes showed very little recruitment compared to SN- and subH-onset dynamotypes (Fig.7a), and recruitment delays for supH-onset dynamotypes were variable (Fig. 7b). When supH-onset dynamotypes did recruit node 2, they did so with less recruitment energy than SN- and subH-onset dynamotypes (Fig.7c). These numerical results align with the theoretical result that supH-onset dynamotypes recruit only through mechanism B, reducing overall recruitment likelihood but also sensitivity to the speed and energy of recruitment. Hence, the model predicts that seizures that start with amplitude scaling are less likely to spread compared to seizures that do not have amplitude scaling at onset.

**Fig 7.**
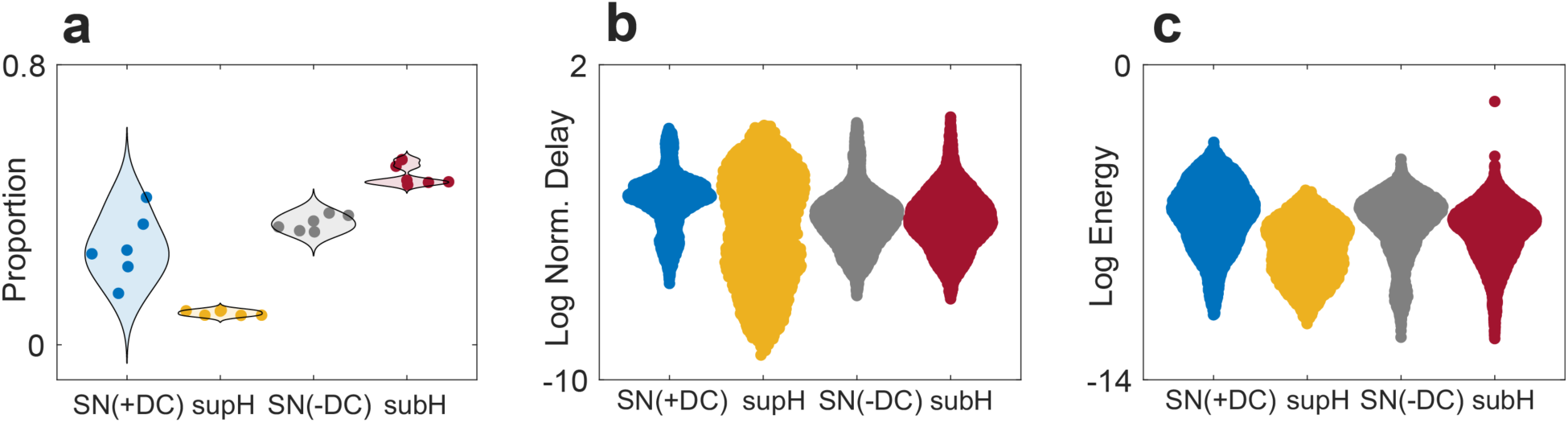
**(a)** Recruitment proportion by node 1 onset. Each datapoint represents one node 1 bursting path. **(b)** Log of normalized recruitment delay and **(c)** log of recruitment energy by node 1 onset. Each datapoint represents one simulation. SN(+DC) = saddle node with baseline shift, supH = supercritical Hopf, SN(-DC) = saddle node without baseline shift, subH = subcritical Hopf.

Recruitment proportions versus coupling strengths and node 1 excitability values are displayed in Fig.8. Across all simulations, the likelihood of recruitment increased with node 1 excitability and with the strength of the coupling between the nodes (Fig.8a). Recruitment patterns varied by onset bifurcation. SN(+DC)-onset dynamotypes showed a gradual increase in recruitment proportion with coupling strength (Fig.8b). In contrast, SN(-DC) and subH-onset node 1 dynamotypes displayed an abrupt increase in recruitment proportion at a coupling strength of 0.02, with a smaller dependence on node 1 excitability (Fig.8c, e). The model therefore predicts that seizures that exhibit baseline shifts or arbitrary amplitude at onset are more likely to recruit when the onset zone is well-connected to potential propagation zones. The excitability of the onset zone has a greater influence on recruitment for seizures that start with baseline shifts compared to those that exhibit arbitrary amplitude at onset.

**Fig 8.**
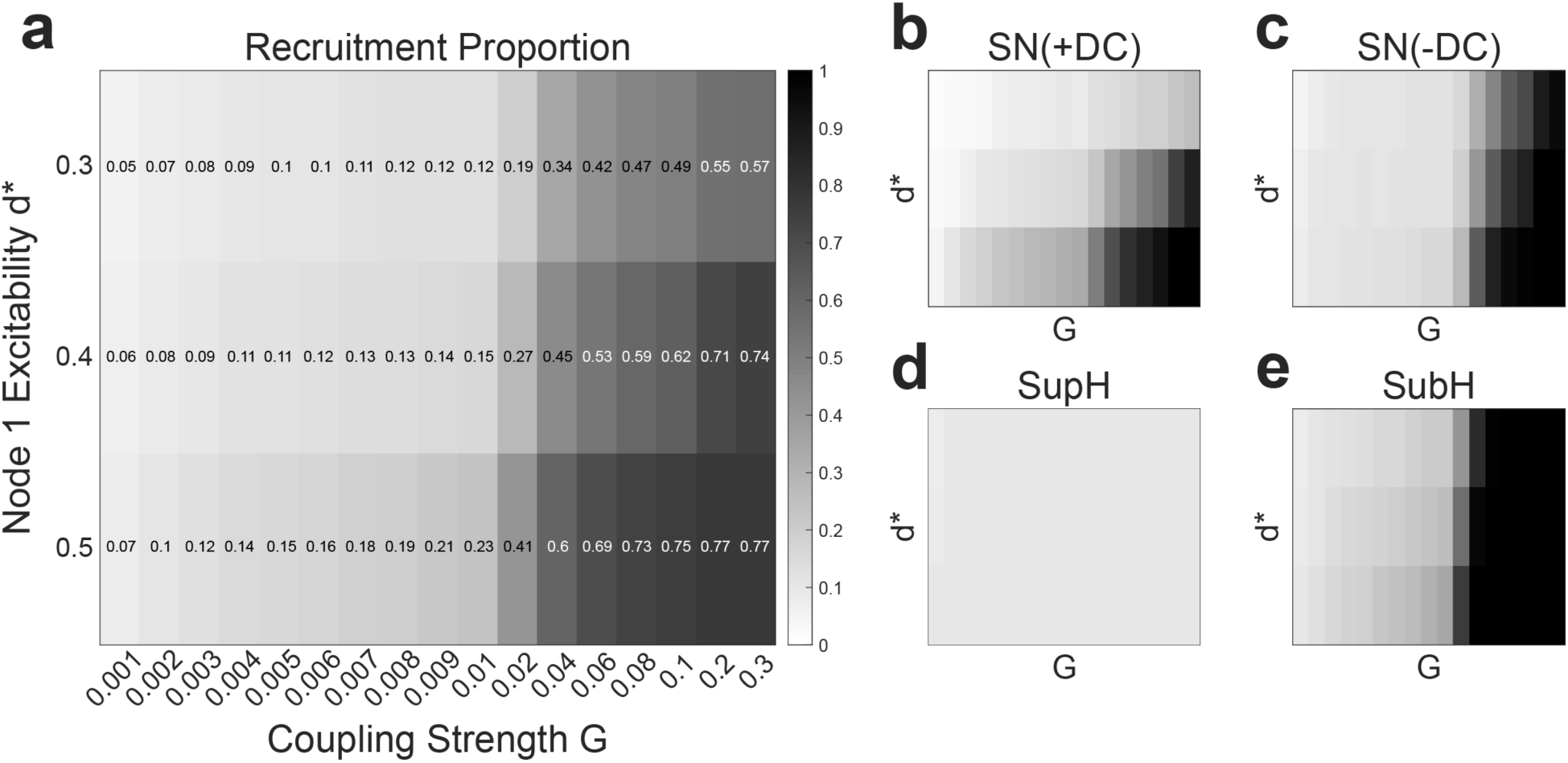
**(a)** Recruitment proportions versus coupling strength (G) and node 1 excitability (d*) for all simulations. Recruitment increased with coupling strength and node 1 excitability. **(b)-(e)** Recruitment proportions versus coupling strength and node 1 excitability for each node 1 onset bifurcation. Dependence on coupling strength and node 1 excitability varied by node 1 onset bifurcation. SN(+DC) = saddle node with baseline shift, supH = supercritical Hopf, SN(-DC) = saddle node without baseline shift, subH = subcritical Hopf.

From theoretical analysis, we predicted that recruitment through mechanism A would be more sensitive to coupling strength and node 1 excitability compared to recruitment through mechanism B. It was no surprise then that supH-onset node 1 dynamotypes, which can only recruit through mechanism B, exhibited a consistent, low recruitment proportion across all coupling strengths and node 1 excitabilities (Fig. 8d). The model predicts these factors do not influence recruitment for seizures that exhibit increasing amplitude at onset.

For consistency with prior work [12,27], we constrained the parameters of the fast subsystem of both nodes to the surface of a sphere centered at the origin. However, only the topology of the bifurcation curves in parameter space is guaranteed, by the fact that the fast subsystem is based on the normal form of the unfolding of a bifurcation, and nothing can be said a priori on their geometry, and thus distances. In the present study, the parameter space geometry influenced the distance between the nodes at the onset of coupling. Thus, to mitigate the possibility that the presented results were a byproduct of this specific parameter space geometry, we confirmed the results were not qualitatively influenced by the initial distance between the nodes. First, we verified that node 2 did not tend to be closer to subH or SN(-DC)-onset node 1 dynamotypes compared to supH and SN(+DC)-onset node 1 dynamotypes (Fig.S3). Then, we analyzed the effect of distance on recruitment properties. There were some relationships between distance and recruitment proportion, delay, and energy, but comparisons between SN(+DC)-, supH-, SN(-DC)-, and subH-onset dynamotypes were generally preserved across distance (Fig.S4-S6).

In the hysteresis-loop implementation, supH-onset dynamotypes cross a SN bifurcation prior to the supH bifurcation. In this work, we adjusted the initial conditions such that the supH-onset dynamotypes did not undergo a SN bifurcation prior to the supH bifurcation. Specifically, we placed the node on the fixed point corresponding to active rest at the start of the simulation. Then, we defined burst onset as the onset of sustained oscillations. In our coupling equations, the coupling term is set to zero until burst onset, meaning that only the supH bifurcation influenced recruitment for supH-onset node 1 dynamotypes. This choice of onset definition facilitated study of the supH-onset bifurcation in isolation. To examine the recruitment properties of the SN(+DC) and supH onset bifurcations in sequence, we re-ran the simulations for supH-onset dynamotypes with a different set of initial conditions, such that a SN bifurcation preceded the supH bifurcation. For these simulations, we defined burst onset as the first departure from rest, i.e. the SN bifurcation. We found that supH-onset node 1 dynamotypes still displayed low recruitment proportion, especially compared to SN(-DC)- and subH-onset node 1 dynamotypes, and variable recruitment delay (Fig.S7). Unsurprisingly, recruitment energy was similar to SN(+DC)-onset node 1 dynamotypes. Recruitment proportion increased with coupling strength, but only for high node 1 excitability (Fig.S8). Thus, the SN(+DC) followed by supH onset bifurcation sequence demonstrated recruitment properties between those of SN(+DC) and supH onset bifurcations in isolation.

Finally, to ensure the results were robust to choice of coupling term (see Discussion), we re-ran the simulations after replacing 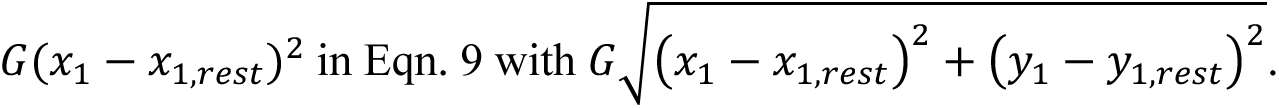 We found the results were indistinguishable (Figs. S9-S10).

#### 3.2.2 Recruitment properties with respect to node 2 initial location

The initial location of node 2 in parameter space also influenced recruitment properties. For most node 2 initial locations, recruitment proportion was consistently low (Fig.9). Recruitment proportions increased and more node 2 initial locations were recruited at high excitability and coupling strength values (Fig.S11). Generally, recruitment delays and energy were consistent and decreased moving from near the LCs region to near the LCb region (Fig.9). These results are consistent with recruitment via mechanism A. Recall that in mechanism A, node 2 reaches node 1 in parameter space before node 1 leaves its onset bifurcation. Thus, recruitment through mechanism A is likely to be low and increase with node 1 excitability and coupling strength. Further, recruitment delay and energy is stereotyped because node 2 can only be recruited in a narrow timespan.

**Fig 9.**
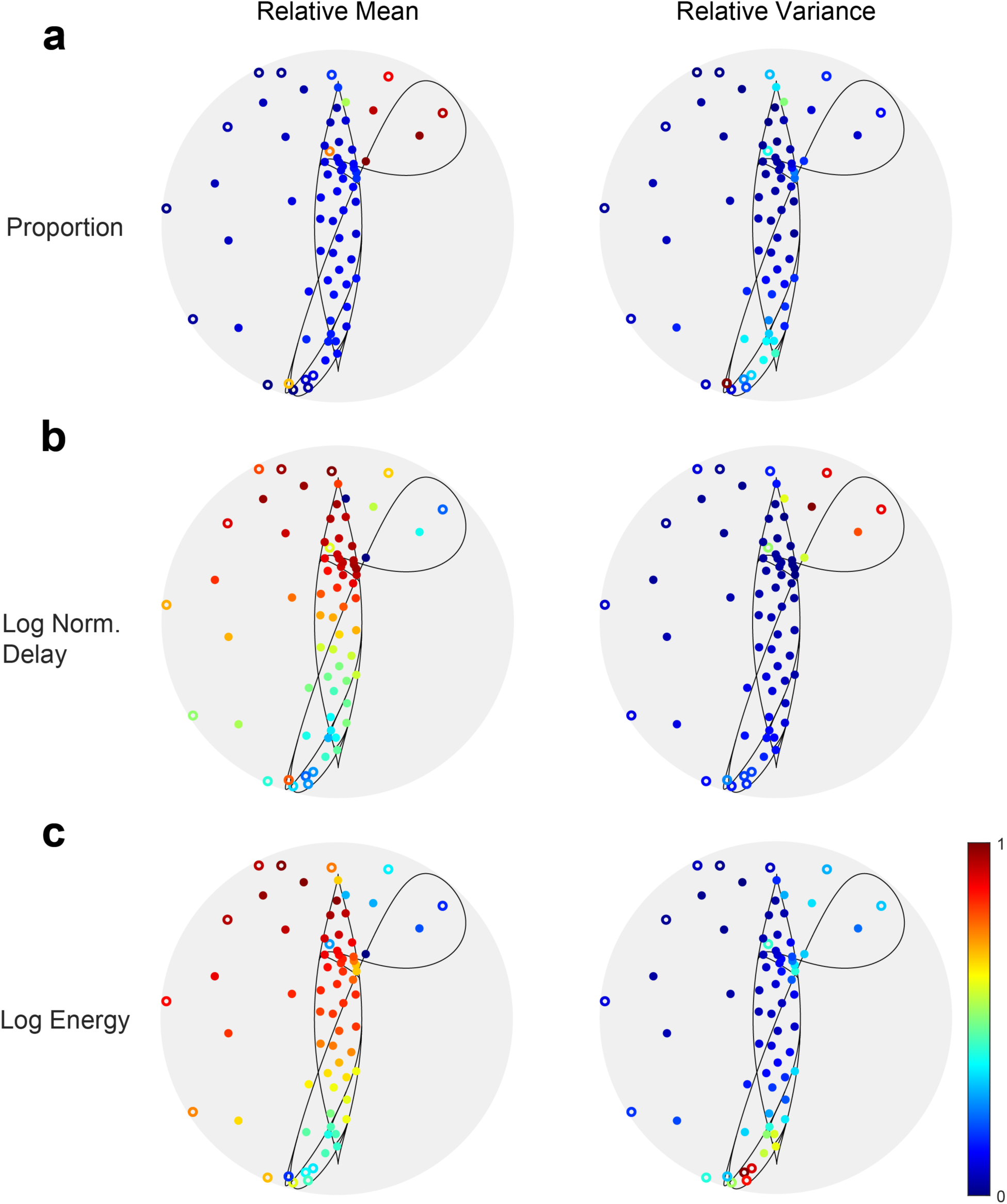
Relative mean and variance of **(a)** recruitment proportion, **(b)** log of normalized recruitment delay, and **(c)** log of recruitment energy by node 2 initial location. For most points, recruitment proportion was consistently low.

A set of initial locations in or near active rest – in the rest region, in the upper right quadrant of the sphere, very close to or to the right of the SN bifurcation – departed from the general trends. These initial locations demonstrated elevated recruitment proportions (Fig. 9). They tended to be recruitable regardless of node 1’s onset bifurcation, even at low node 1 excitability and low coupling strength (Fig.S11). They were the only initial locations recruited by supH-onset node 1 dynamotypes (Fig.S11). These observations are consistent with the theoretical prediction that nodes in active rest can easily be recruited via mechanism B. Entering either the LCs region or the ictal-only region on the path to LCb facilitated recruitment of nodes in active rest. Node 1 failed to recruit node 2 from active rest only if node 2 did not move into the LCs bistability or ictal-only regions, which occurred only for simulations with sufficiently low coupling strength and node 1 excitability. Initial locations in or near active rest also showed lower and more variable recruitment delays, and reduced recruitment energy (Fig. 9). The proximity of these nodes to the LCs and ictal-only regions likely lowered recruitment delays and energy. Recruitment through mechanism B can occur at any point in the node 1 burst, facilitating variation in recruitment delay.

These results illustrate that the dynamical properties of the potential propagation zone influence its likelihood and speed of recruitment. For most initial dynamics, the model suggests propagation zones are recruited infrequently and with stereotypical delay. However, nodes that are initially in active rest are more likely to be recruited and demonstrate more variable recruitment delays.

Normalized recruitment delay and recruitment energy increased as the initial location of node 2 moved upwards on the sphere. A set of initial locations in the rest region in the top half of the sphere departed from these trends, instead showing more frequent recruitment, more variable recruitment delay, and lower and less variable recruitment energy.

#### 3.2.3 Propagation zone dynamics *in silico*

Although not mandatory, clinicians expect that recruited tissue most commonly will adopt the same bursting dynamics as the onset zone. Prior computational models of coupled nodes studying seizure propagation typically implement this expectation by imposing the same dynamics on all nodes. Here we did not impose this constraint, instead allowing seizure focus and propagation nodes to have different intrinsic dynamics. The coupling we designed facilitated the occurrence of similar dynamics, but also allowed for situations where that may not be true. Specifically, recruitment through mechanism B could lead to supH onset dynamics in node 2 even if node 1 was not supH-onset. Further, we found that node 2 could experience a drift in the location of the resting state while traversing parameter space, which appeared similar to a baseline shift in the timeseries. Note that we use the term *drift* rather than *shift* to distinguish these changes in baseline from the abrupt changes at burst onset caused by a SN bifurcation. Breaking of timescale separation in some simulations with sufficiently high coupling strength also allowed the onset pattern of node 2 to vary from that of node 1. Figs. 10-12 demonstrate the range of node 2 dynamics possible for each node 1 onset. For this analysis, we grouped SN(-DC) and subH onsets together because these dynamotypes are indistinguishable in timeseries data [24].

**Fig 10.**
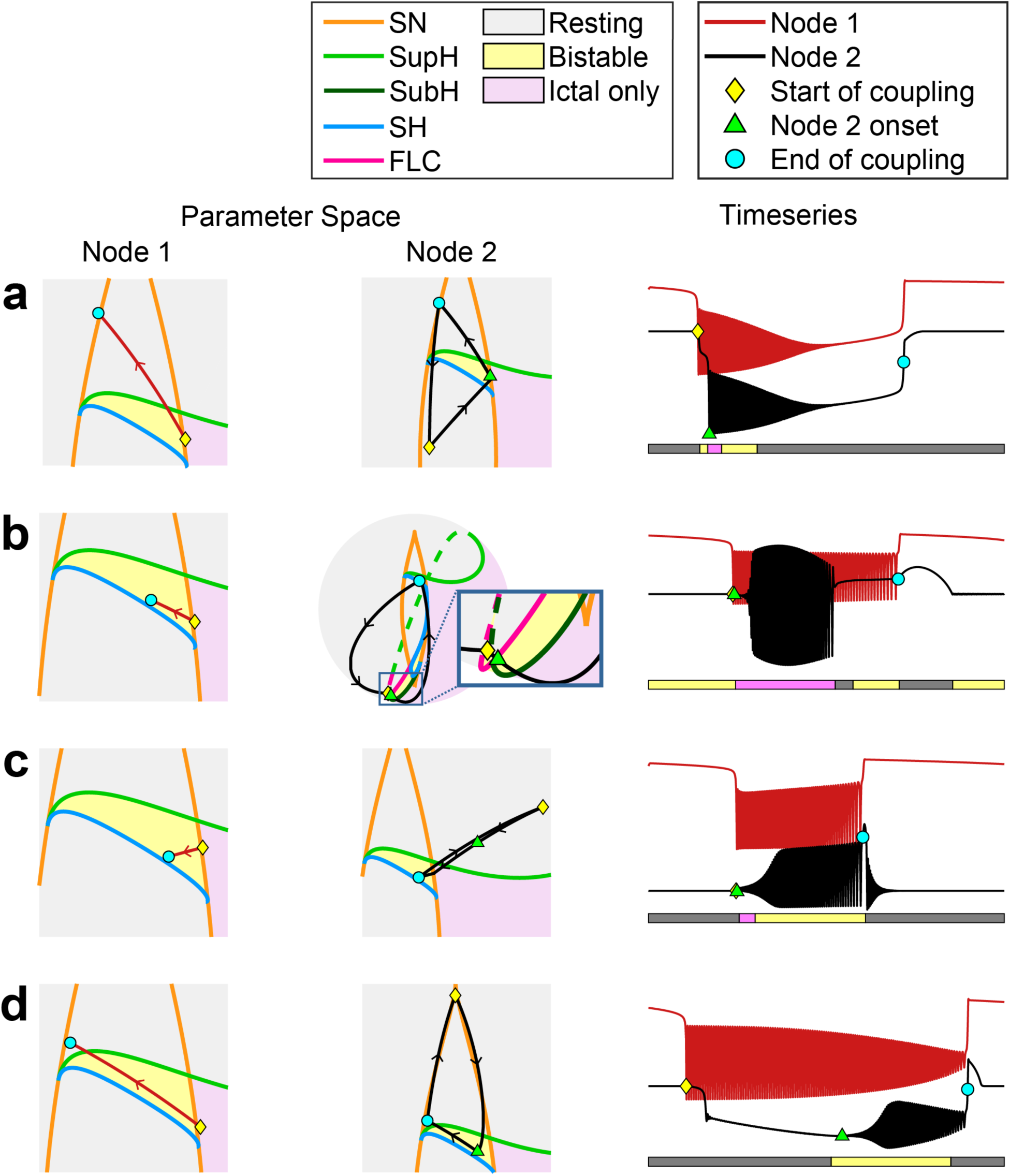
Recruitment examples for SN(+DC)-onset node 1 dynamotypes. Parameter space movement is shown on the left. Timeseries (offset for visualization) are shown on the right. The shaded bars under the timeseries indicate the parameter space region of node 2. Arrows indicate the direction of travel in parameter space. Yellow diamonds mark the location of node 1 and node 2 in parameter space at the time of coupling onset. Cyan circles mark the location of node 1 and node 2 in parameter space at the time of coupling offset. Green triangles mark the location of node 2 in parameter space at node 2 onset. The symbols are also shown in the node 2 timeseries (right). **(a)** Example in which the onset dynamics of node 2 matched that of node 1. **(b)-(d)** Examples in which the onset dynamics of node 2 differed from that of node 1. In (b), the box zooms into the LCb region for node 2. SN(+DC) = saddle node with baseline shift, SN = saddle node, SupH = supercritical Hopf, SubH = subcritical Hopf, SH = saddle homoclinic, FLC = fold limit cycle.

**Fig 11.**
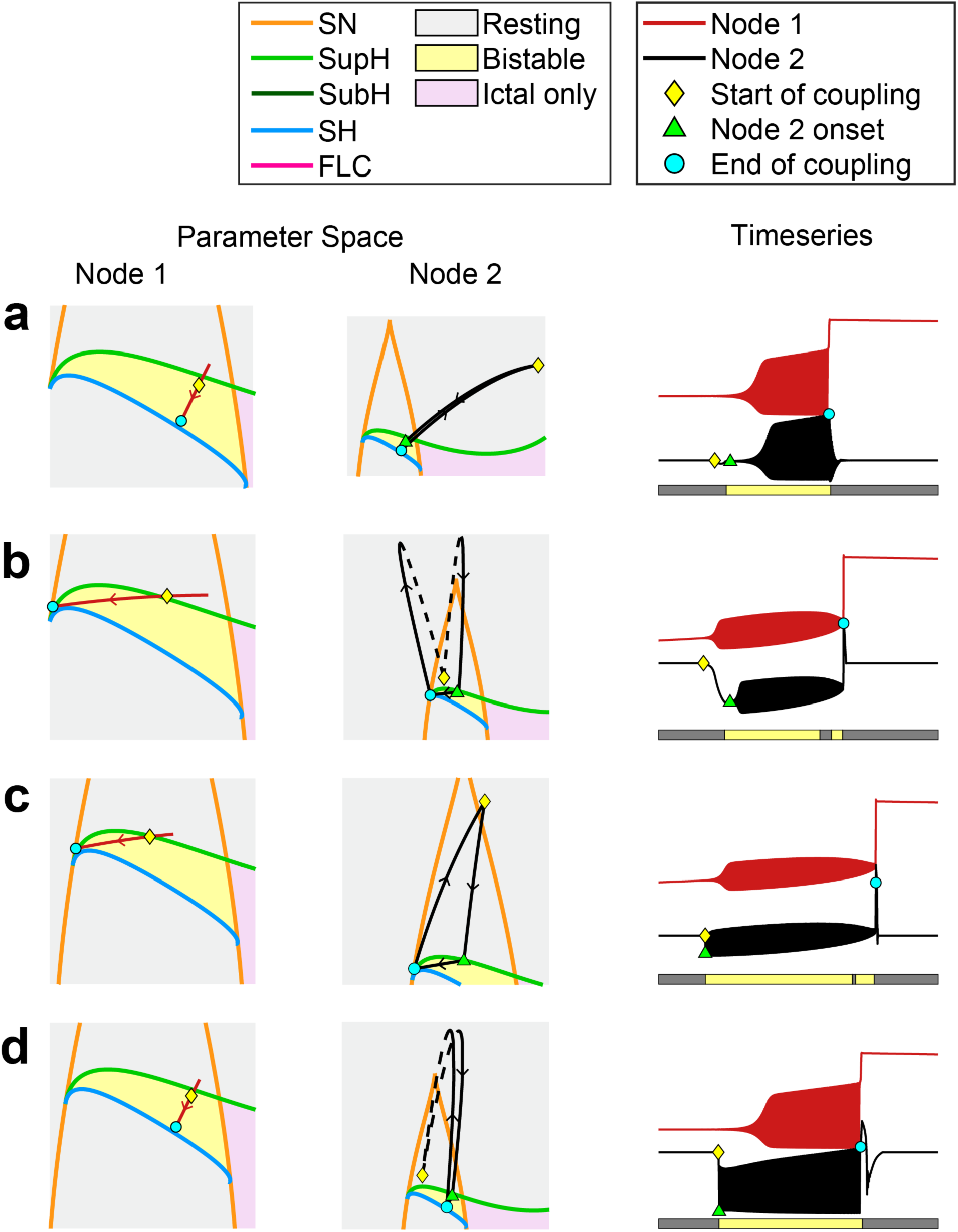
Recruitment examples for supH-onset node 1 dynamotypes. Parameter space movement is shown on the left. Timeseries (offset for visualization) are shown on the right. The shaded bars under the timeseries indicate the parameter space region of node 2. Arrows indicate the direction of travel in parameter space. Yellow diamonds mark the location of node 1 and node 2 in parameter space at the time of coupling onset. Cyan circles mark the location of node 1 and node 2 in parameter space at the time of coupling offset. Green triangles mark the location of node 2 in parameter space at node 2 onset. The symbols are also shown in the node 2 timeseries (right). In **(a)** and **(b)**, node 2 exhibited increasing amplitude from zero at onset. In **(c)** and **(d)**, node 2 exhibited arbitrary amplitude at onset. In (b) and (d), there was also a baseline drift at or just before onset. SN = saddle node, SupH = supercritical Hopf, SubH = subcritical Hopf, SH = saddle homoclinic, FLC = fold limit cycle.

**Fig 12.**
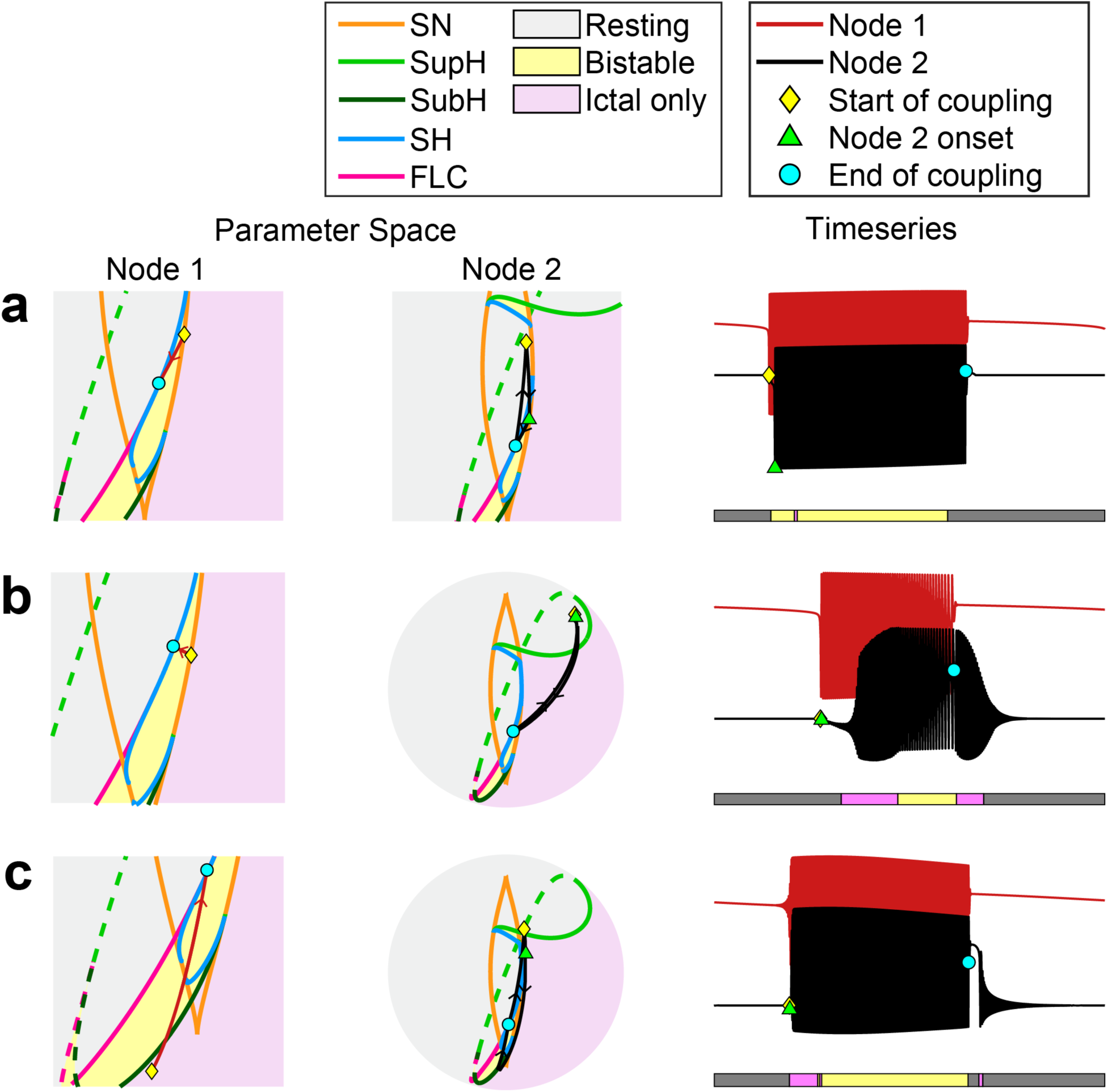
Recruitment examples for subH- or SN(-DC)-onset node 1 dynamotypes. Parameter space movement is shown on the left. Timeseries (offset for visualization) are shown on the right. The shaded bars under the timeseries indicate the parameter space region of node 2. Arrows indicate the direction of travel in parameter space. Yellow diamonds mark the location of node 1 and node 2 in parameter space at the time of coupling onset. Cyan circles mark the location of node 1 and node 2 in parameter space at the time of coupling offset. Green triangles mark the location of node 2 in parameter space at node 2 onset. The symbols are also shown in the node 2 timeseries (right). In **(a)**, node 2 exhibited arbitrary amplitude at onset. In **(b)**, node 2 displayed increasing amplitude at onset. In **(c)**, node 2 showed arbitrary amplitude at onset and a drift in baseline during the burst. SN(-DC) = saddle node without baseline shift, SN = saddle node, SupH = supercritical Hopf, SubH = subcritical Hopf, SH = saddle homoclinic, FLC = fold limit cycle.

SN(+DC)-onset node 1 dynamotypes occur in the LCs region and display a baseline shift and arbitrary amplitude at onset. For most initial locations recruited by SN(+DC)-onset dynamotypes, recruitment occurred through mechanism A and the onset dynamics of node 2 matched those of node 1, as in Fig.10a. For some initial locations, recruitment occurred through mechanism B and distinct onset dynamics were observed in node 2. When node 2 was pulled from near the LCb region across the ictal-only region of the sphere, the burst in node 2 appeared unrelated to that in node 1 (Fig.10b). Node 2 crossed a subH bifurcation to enter the ictal-only region and thus did not demonstrate a baseline shift at onset. The burst in node 2 also terminated before the burst in node 1 since node 2 crossed an offset bifurcation upon leaving the ictal-only region. As demonstrated in Fig.10c-d, node 2 could also demonstrate increasing amplitude at onset when node 1 pulled it across the supH bifurcation to enter LCs. If node 2 crossed from rest to active rest prior to entering the LCs region, a baseline drift could precede burst onset in node 2 (Fig.10d).

SupH-onset node 1 dynamotypes occur in the LCs region and exhibit increasing amplitude from zero at onset. SupH-onset node 1 dynamotypes only successfully recruited through mechanism B and when node 2 was in or near active rest (Fig.S11). Typically, recruited node 2’s displayed increasing amplitude from zero at onset, also characteristic of supH (Fig.11a-b). However, if node 2 moved quickly into the LCs region, onset lacked amplitude scaling (Fig.11c-d). A baseline drift could precede burst onset if node 2 crossed from rest to active rest prior to entering the LCs region (Fig.11b, d).

SN(-DC)- and subH-onset node 1 dynamotypes occur in the LCb region and exhibit arbitrary amplitude at onset without baseline shift. Nodes were often recruited by SN(-DC)- or subH-onset node 1 dynamotypes through mechanism A, resulting in the same onset dynamics, as in Fig.12a. If node 2 was initially located in or near active rest, then coupling dragged the node across the supH bifurcation into the ictal-only region of the sphere, resulting in recruitment through mechanism B. If movement across the supH bifurcation was sufficiently slow, node 2 displayed amplitude scaling at onset (Fig.12b). If node 2 moved quickly across the supH and subH bifurcations, supH onset characteristics could be lost (Fig.12c).

Further, baseline drifts smaller than the amplitude of the oscillations were possible in node 2 (Fig.12c).

To summarize, the model predicts that seizure onset dynamics can remain the same or change during propagation from the seizure onset zone to propagation zone. Mechanism A promotes similar dynamics, while mechanism B tends to facilitate supH or, to a smaller degree, subH onset dynamics, which may be distinct from the onset dynamics of node 1. Further, simulations revealed that baseline drifts could occur as node 2 traversed parameter space, and sufficiently high coupling strength could break timescale separation, leading to unexpected node 2 dynamics. Together, these effects allow for changes in baseline and amplitude scaling at onset to occur in seizure onset zone only, propagation zone only, neither, or both.

### 3.3 Propagation zone dynamics in clinical data

Several of the simulations produced mismatched seizure onset zone and propagation zone dynamics, an unexpected situation clinically. We therefore searched for examples of the combinations of onset dynamics demonstrated in Figs. 10-12 in human intracranial EEG data. We identified recordings compatible with specific onset bifurcations visually: SN(+DC) onsets were identified via baseline shifts, supH onsets were identified via increasing amplitude, and SN(-DC) or subH onsets were identified via arbitrary amplitude. It needs to be stressed that such analysis of shifts and amplitude scaling laws does not prove that a bifurcation is occurring, but only that the data are compatible with this scenario [23]. For consistency with the definition of burst onset used in our simulations, we specifically considered the dynamics in the onset channel at the onset of *sustained* oscillations.

The model predicted that, when recruited by SN(+DC)-onset node 1 dynamotypes, node 2 could demonstrate amplitude scaling or arbitrary amplitude, with or without a change in baseline (Fig.10). The clinical example displayed in Fig.13 confirms this range of propagation zone dynamics is possible in humans. This seizure began in the basal temporal region with a clear baseline shift at onset, compatible with a SN(+DC) bifurcation. The propagation zone included the right parietal, lateral temporal, and anterolateral basal temporal regions, which demonstrated dynamics consistent with subH or SN(-DC), SN(+DC), and supH onset bifurcations, respectively. The anterolateral basal temporal region also demonstrated a change in baseline prior to amplitude scaling, which could be due to a baseline drift preceding the supH onset bifurcation as in Fig. 10d. In the context of coupled nodes of the Multiclass Epileptor, these differing onset patterns suggest the initial dynamical states of the propagation channels varied. Fig.S12 provides another example of possible SN(+DC) onset in the onset zone, followed by a possible supH onset in the propagation zone.

**Fig 13.**
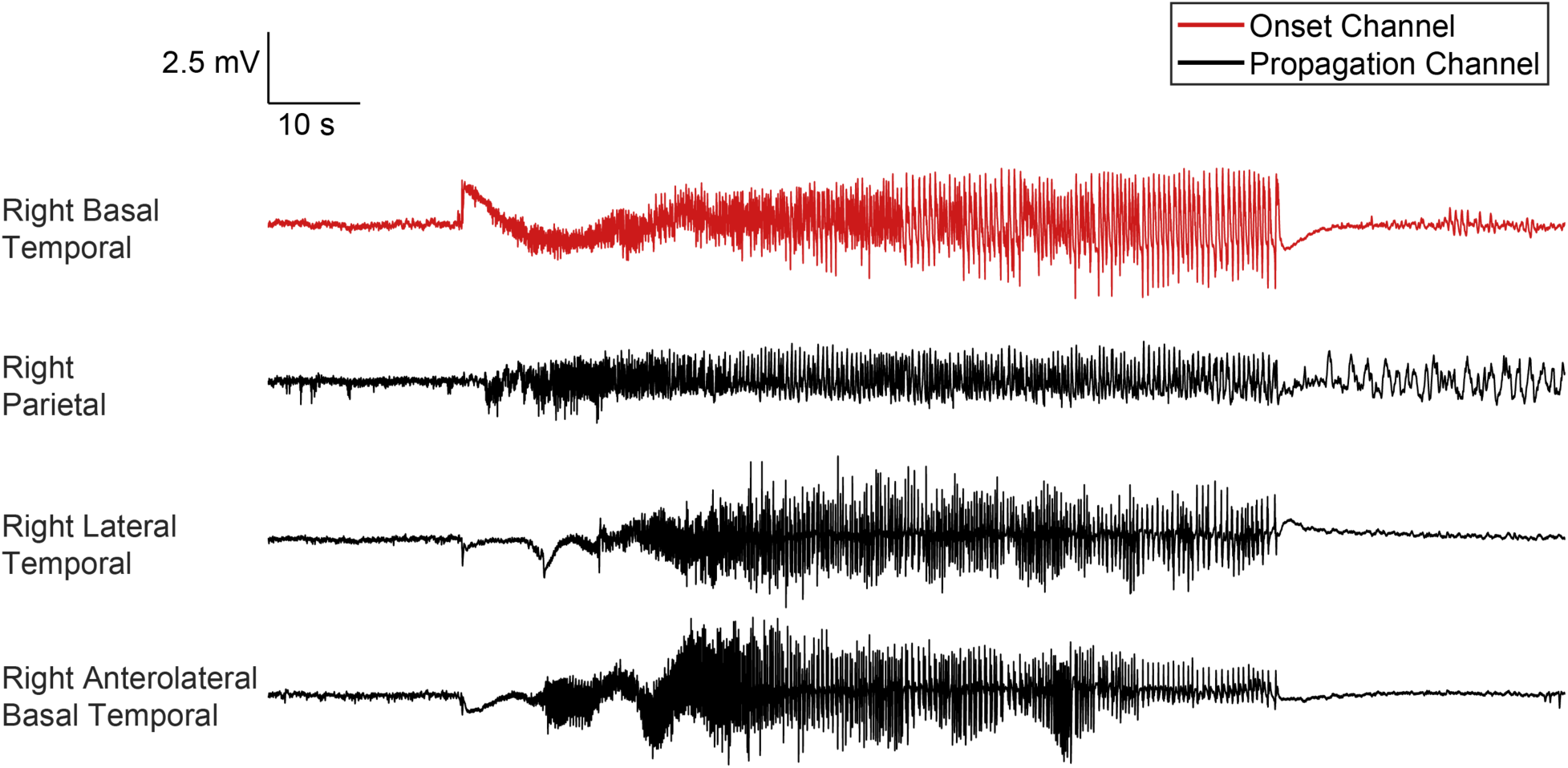
Clinical seizure propagation example 1. There is a baseline shift at onset in the onset channel, compatible with a SN(+DC) onset bifurcation. Propagation channel dynamics vary. The right parietal channel exhibits arbitrary amplitude without a change in baseline at onset, consistent with a subH bifurcation. The right lateral temporal channel shows a baseline shift at onset, consistent with a SN(+DC) bifurcation. The right anterolateral basal temporal channel displays a baseline shift followed by increasing amplitude at onset, consistent with a baseline drift followed by a supH bifurcation.

In the model, node 2 could demonstrate amplitude scaling or arbitrary amplitude, with or without baseline drift, at onset when recruited by supH-onset node 1 dynamotypes (Fig.11). In clinical data, we found examples of amplitude scaling, changes in baseline, and arbitrary amplitude at onset in the propagation zone. The seizure in Fig.S13 demonstrated increasing amplitude at onset – indicative of a supH bifurcation – in both the seizure onset zone and propagation zone. In Fig.14, the seizure began in the left hippocampus with increasing amplitude and without a baseline shift at onset. After a delay, the seizure spread to propagation channels in the right hippocampus, where there was a change in baseline at onset. In Fig.15, the onset channel demonstrates amplitude scaling at the onset of sustained oscillations. The propagation channels demonstrated arbitrary amplitude at onset and appeared to fire prior to the onset of sustained oscillations in the onset channel. In coupled nodes of the Multiclass Epileptor, we observed similar onset behavior when coupling was strong (Fig.11c).

**Fig 14.**
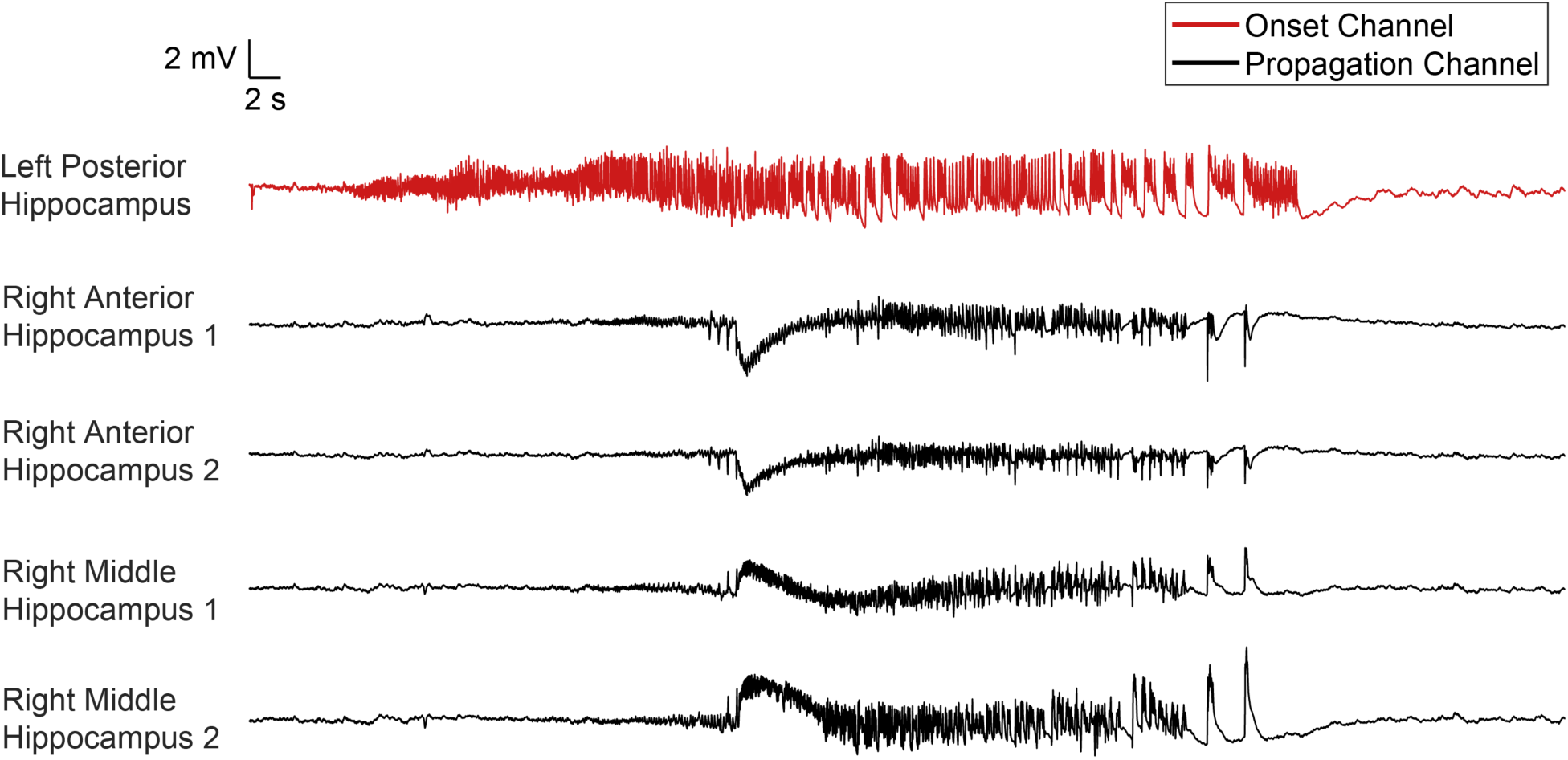
Clinical seizure propagation example 2. There is increasing amplitude at onset in the onset channel, compatible with a supH bifurcation. The propagation channels display clear changes in baseline at onset. Numbers indicate electrode contacts.

**Fig 15.**
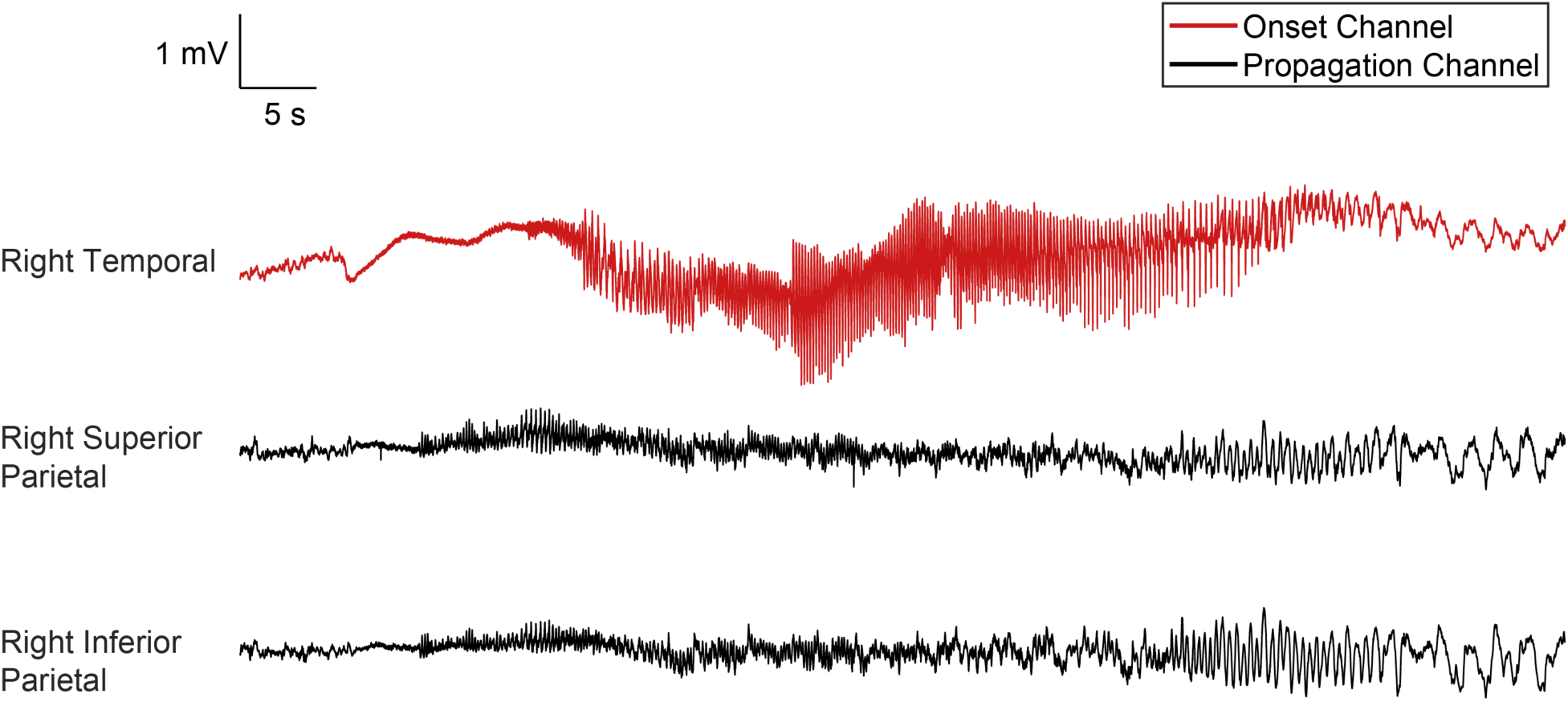
Clinical seizure propagation example 3. Increasing amplitude at onset of sustained oscillations in the onset channel is compatible with a supH bifurcation. The propagation channels display arbitrary amplitude at onset.

When node 1 was a subH- or SN(-DC)-onset dynamotype, node 2 demonstrated amplitude scaling or arbitrary amplitude at onset in the model (Fig.12). Baseline drifts could also occur in node 2. We found a similar range of behavior in clinical data. The seizure in Fig.16 began with arbitrary amplitude in the seizure onset zone, suggesting a SN(-DC) or subH onset. In the propagation zone, there was increasing amplitude at onset, implying a supH bifurcation. There also may have been a baseline drift >5 sec prior to the supH bursting. We did not witness an example of a baseline drift temporally separated from supH onset dynamics in node 2 when node 1 was subH- or SN(-DC)-onset in the model. However, node 2 could display this behavior if it began in rest was pulled to active rest and then into the ictal-only region across the supH bifurcation. This path did not occur in the simulations presented here because node 2 moved along the shortest path to node 1 at each point in time. A slightly longer path could produce the observed sequence. Fig.S14 confirms the potential for subH- or SN(-DC)-like dynamics (arbitrary amplitude) at onset in both the seizure onset zone and propagation zone.

**Fig 16.**
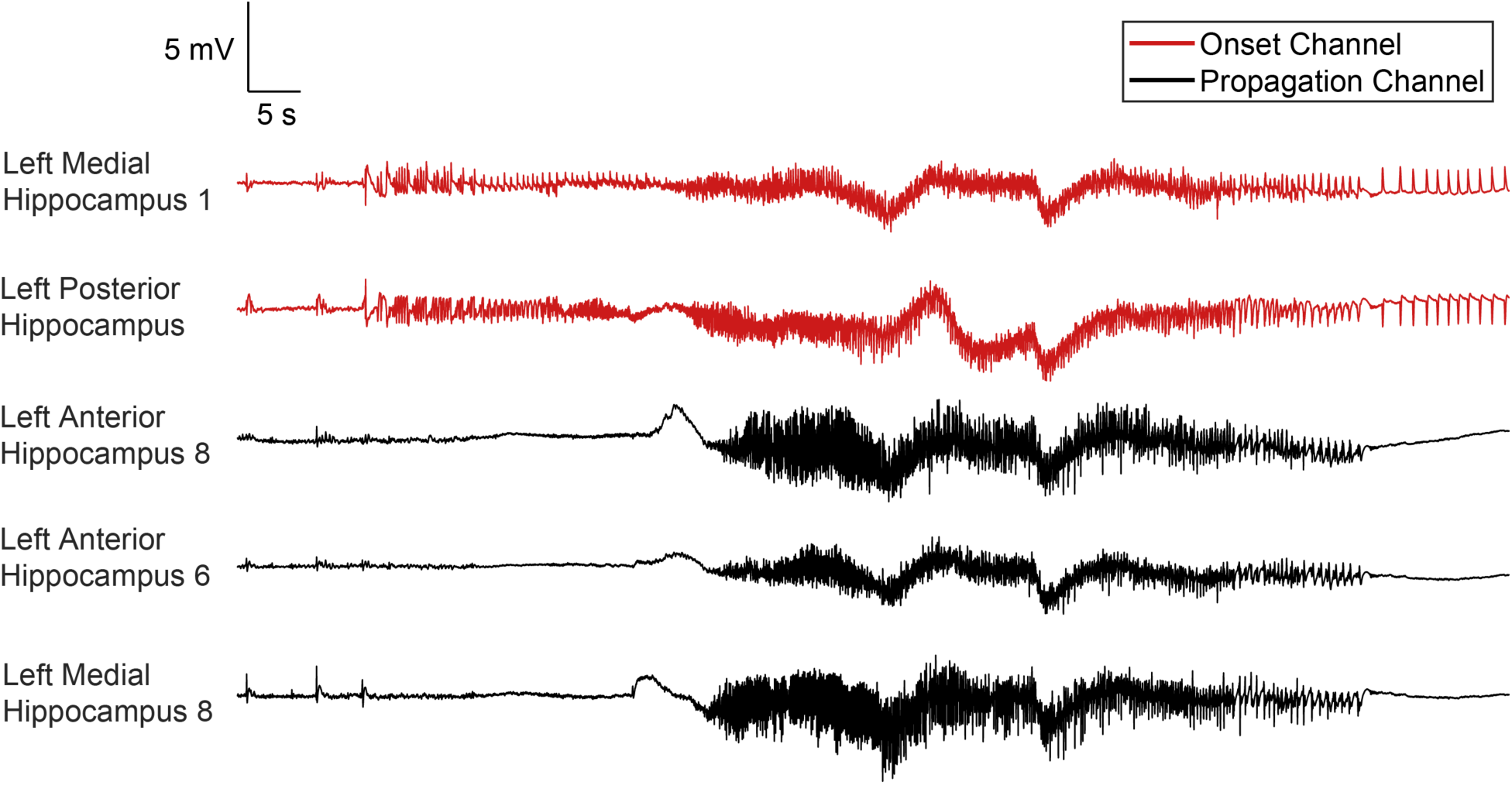
Clinical seizure propagation example 4. The onset channels display arbitrary amplitude at onset, compatible with a subH or SN(-DC) onset bifurcation. The propagation channels display increasing amplitude at onset, suggesting a supH onset bifurcation. There is also potentially a baseline shift without bursting before the supH onset. Numbers indicate electrode contacts.

## 4. Discussion

### 4.1 Factors impacting recruitment

We evaluated the influence of onset zone excitability, coupling strength, onset zone dynamotype, and propagation zone dynamics on recruitment likelihood, delay, and energy. The model predicted that recruitment properties depend on seizure onset zone dynamotype. Specifically, seizures that exhibited arbitrary amplitude without a baseline shift at onset recruited more often and faster than seizures that exhibited a baseline shift at onset. Seizures that displayed increasing amplitude from zero at onset were unlikely to propagate compared to seizures with arbitrary amplitude at onset. While quantitative validation of model predictions is beyond the scope of this study, it is worth noting that generalized seizures – which propagate quickly and extensively – tend to exhibit arbitrary amplitude without a baseline shift at onset. In contrast, baseline shifts at seizure onset are a strong indicator of an isolated focal epilepsy [32–34], in which seizures do not always propagate quickly or extensively and which responds well to focal resective surgery. Finally, the initial dynamics of the potential propagation zone influenced its likelihood of recruitment. In other words, some resting states were predicted to be more vulnerable to recruitment than others. These predictions provide a guide for future experiments to improve the methods to quantify and control human seizure spread.

### 4.2 Choice of coupling method

A unique aspect of our model was the ability to simulate the effects of a wide range of potential dynamics in both the seizure onset zone and in the propagation zone. We simulated eight seizure onset zone dynamotypes and 71 initial propagation zone dynamics, including both bistable and monostable resting states. In contrast, previous dynamical models of epilepsy networks have focused on single dynamotypes and generally require bistability [14,19–21,35,36]. Strategies for connecting nodes within these models have included fast-to-fast coupling [14,35,36] and fast-to-slow coupling [19–21]. Fast-to-fast coupling aims to mimic the influence of white matter connections, which may be inferred from EEG or MRI data in the context of patient-specific modeling. Fast-to-slow coupling instead models the influence of fast variables (e.g. neural firing) in one brain region on slow variables (e.g. extracellular ion concentrations) in another brain region. Compared to fast-to-fast coupling, fast-to-slow coupling may better account for long recruitment times (up to seconds) during seizure propagation. Fast-to-slow is the coupling method utilized to connect Epileptors, from which we drew inspiration in the present work.

To investigate how a rich array of seizure onset zone and propagation zone dynamics influence recruitment, we introduced a novel method to connect two nodes in the Multiclass Epileptor. While different onset/offset bifurcation curves can be obtained by tuning the parameters of well-known models such as the Jansen-Rit [37] and its extension, the Wendling-Chauvel [38], these models typically lack slow dynamics for autonomous seizure generation. To our knowledge, the Multiclass Epileptor is the only model in the literature that can reproduce a variety of dynamotypes (i.e. a range of bifurcation onsets and offsets) seen in human seizures in an easily controllable way. Different dynamotypes are realized by navigating the parameter space via the slow subsystem of the model. To connect two nodes of the model, we implemented a novel form of fast-to-slow coupling, allowing fast activity in node 1 (the seizure onset zone) to alter the dynamics of node 2 (the propagation zone). Specifically, the fast subsystem of node 1 determined how quickly the parameters of node 2 (i.e. its slow subsystem) changed. The location of node 1 determined the direction of movement of node 2. Based on the clinical observation that the dynamics of the propagation zone are likely to match those of the seizure onset zone, we assumed that the parameters of node 2 tend to become similar to those of node 1 during recruitment. This assumption also facilitates synchronization, a phenomenon commonly observed in clinical EEG. Under this assumption, we could place node 2 in rest-only regions of parameter space, facilitating a focus on recruitment of brain regions incapable of independent seizure activity. Because the model’s parameters are phenomenological and lack an explicit biophysical interpretation which would constrain the choice of physiological parameter values for the rest location, we systematically explored all possibilities.

A key aspect of the Multiclass Epileptor is timescale separation between the fast and slow subsystems. Timescale separation, guaranteed by setting 𝑘_1_ << 1 in Eqn. 5, follows from the progression toward seizure termination, over the course of seconds or minutes, as is seen in human epilepsy, being much slower than the oscillations during the ictal phase [39]. However, timescale separation did not hold in all examples of our model of seizure propagation. Within our propagation model, timescale separation is broken for node 2 when coupling strength is sufficiently high. Breaking of timescale separation enables recruitment of node 2 from regions of parameter space that are far from bifurcations and from node 1. The mathematical implication of breaking timescale separation is that the topology of node 2’s state space changes faster than node 2’s fast subsystem can converge to a stable solution, and also that the parameter space map itself is not guaranteed to be correct. As such, node 2’s EEG (x variable) may show unexpected behavior; for example, node 2 may cross a supH bifurcation but lack amplitude scaling (as in Fig.11c) if the movement on the map is sufficiently fast. There is physiological rational for this situation. When a clinical seizure begins in the seizure onset zone, the propagation zone is recruited quickly – often within milliseconds and rarely after more than a few seconds [40]. That recruitment can span the entirety of the brain, and it is unlikely every brain node is always at the same dynamical baseline. The example in Fig. 14, where one dynamotype begins on the left, then triggers a different one on the right, corroborates this idea. Thus, we conclude that our method of recruitment is plausible.

While rigorous validation of the model is beyond the scope of this work, we compared to EEG recordings in patients with epilepsy to ensure recruitment behaviors predicted by the model were possible clinically. The model allowed for conditions in which seizure onset dynamics in the propagation zone did not have to be identical to those of the seizure onset zone. These predictions were unexpected, but we were able to find several examples of similar behaviors in human EEG recordings. One particularly unexpected prediction, from the clinical perspective, was the possibility to observe a change in baseline at onset in the propagation zone but not the seizure onset zone (Fig.11b, d). In focal epilepsy, a DC shift in the propagation zone only is a surprise – baseline shifts have been proposed as a biomarker of the seizure onset zone [32–34]. Yet, we found examples in intracranial EEG from several patients that confirmed this and other predictions (Figs. 13-16, S12-S14).

There are other methods of modeling propagation that we did not perform. For example, node 1 could modulate the excitability of node 2 only and not affect its position on the map, as in coupled Epileptors. This form of coupling would require node 2 to follow a predetermined bursting path in a bistability region of parameter space. By modulating node 2’s excitability, node 1 would influence how quickly node 2 moved along its bursting path (i.e. the speed of node 2’s slow subsystem). However, the choice of bursting path would prescribe node 2’s dynamotype, as well as whether the nodes are capable of synchronicity. Thus, that method would not account for the tendency for the propagation zone to adopt the same dynamical characteristics as and synchronize with the seizure onset zone, if the paths for the two nodes were not identical. The different coupling methods in the literature, fast-to-fast, fast-to-slow on identical paths, and the fast-to-slow on different paths we investigated here, are not mutually exclusive. Future work will investigate their interactions.

There are also other coupling terms that could be implemented. We utilized the novel activity in node 1’s x variable to modulate the speed of node 2’s movement in parameter space (Eqn. 9; speed proportional to (𝑥_1_-𝑥_1,rest_)^2^ during coupling). Since the x variable mimics a voltage recording, our coupling influenced the activity in the potential propagation zone (node 2) based on the change in voltage in the seizure onset zone (node 1). Another option would be to include the change in both fast variables in the coupling term. We repeated the analysis with a modified coupling term including both x and y terms and found the results to be indistinguishable (Figs. S9-S10).

### 4.3 Clinical relevance

#### 4.3.1 Motivation for identifying seizure dynamotypes

Clinical seizure classification is primarily based on semiology and visual analysis of EEG [3,9]. Seizures are first divided into generalized, focal, and unknown onset, and may be subdivided by clinical symptoms and known or suspected etiology. While these are useful descriptive tools, there are little data on how such classifications affect response to treatment [41]. They also ignore the underlying dynamics that define the seizure itself. The dynamotype is a complementary method of seizure classification based upon invariant seizure properties. Dynamical systems theory predicts that different onset bifurcations have distinct properties, which have only recently been described in human epilepsy [24]. Previous work predicted that different dynamotypes are associated with differential response to aborting stimulation [27]. Here, we present the first demonstration of potential differences in seizure propagation among dynamotypes, further underscoring the utility in identifying patients’ dynamotypes. The dynamotype classification provides a language and basis for future studies investigating seizure propagation. That is not to exclude the relevance of other seizure dynamics – onset and offset bifurcation are but two possibly relevant dynamical features. The relevance of different dynamical features to seizures depends on the specific scientific or clinical question that is asked. Here, we chose to focus on onset bifurcation since this is the most relevant to recruitment.

#### 4.3.2. Explanation of unusual propagation zone dynamics

There is very little prior literature analyzing how dynamics change during propagation in human seizures. The clinical expectation is that recruited nodes will adopt the same dynamics as the seizure onset zone. When recruited regions have different dynamics, this situation is often assumed to mean that the second node is an independent seizure focus. However, the results in Figs. 10-12 suggest that a recruited node can adopt distinct dynamics simply due to its resting state position. This prediction provides a novel perspective on seemingly independent propagation zone dynamics.

#### 4.3.2 Toward improved analysis of seizure networks

A contributing factor to poor outcomes in resective surgery for drug-resistant focal epilepsy is a lack of understanding of patients’ seizure networks. A novel method used to better delineate onset and propagation zones is the use of the virtual epileptic patient (VEP), which seeks to recreate patients’ seizure networks in a computational model based on electrographic and imaging data [19,21]. Thus far, nodes in VEP models have been populated with Epileptors, which model the SN(+DC)-SH dynamotype. In demonstrating that seizure recruitment properties depend upon dynamotype, we motivate the inclusion of a range of dynamotypes in future whole-brain network dynamical models of epilepsy. Given that patients’ exhibit different dynamotypes that can change over time [24,26], inclusion of a wider range of dynamics in large scale dynamical models may provide more accurate and patient-specific results.

## Supplemental Information

**Supplementary Table 1:** Node 1 offset 𝑨, onset 𝑩, speed 𝑘_1_, and initial conditions [𝑥_0_, 𝑦_0_, 𝑧_0_].

| Node 1 Class | **A** | | | **B** | | | $k_1$ | $x_0$ | $y_0$ | $z_0$ |
| --- | --- | --- | --- | --- | --- | --- | --- | --- | --- | --- |
| SN(+DC)/SH | 0.3484 | 0.037 | 0.1931 | 0.337 | 0.0753 | 0.2019 | 0.00025 | 0.40 | 0 | 0.07 |
|  | 0.3296 | -0.0243 | 0.2253 | 0.3479 | 0.079 | 0.181 | 0.0005 | 0.50 | 0 | 0.20 |
|  | 0.3448 | 0.02285 | 0.2014 | 0.3426 | 0.0772 | 0.1915 | 0.0005 | 0.40 | 0 | 0.044 |
| SN(+DC)/SupH | 0.3209 | 0.0504 | 0.2353 | 0.3407 | 0.0766 | 0.195 | 0.001 | 0.4026 | 0 | 0.05 |
|  | 0.3102 | -0.0404 | 0.2493 | 0.3496 | 0.0795 | 0.1774 | 0.001 | 0.498 | 0 | 0.267 |
|  | 0.3155 | 0.0017 | 0.2459 | 0.3291 | 0.0727 | 0.2155 | 0.001 | 0.48 | 0 | 0.12 |
| SupH/SH | 0.3464 | 0.0289 | 0.1979 | 0.3216 | 0.0475 | 0.2331 | 0.0001 | -0.63 | 0 | 0.1472 |
|  | 0.3262 | -0.0338 | 0.229 | 0.3175 | 0.0139 | 0.2429 | 7.5e-5 | -0.607 | 0 | 0.1795 |
|  | 0.3517 | 0.0527 | 0.1831 | 0.3222 | 0.0565 | 0.2302 | 0.0001 | -0.633 | 0 | 0.1912 |
| SupH/SupH | 0.3171 | -0.066 | 0.2347 | 0.3177 | 0.0155 | 0.2425 | 2.5e-5 | -0.6311 | 0 | 0.3009 |
|  | 0.3115 | -0.0546 | 0.245 | 0.3129 | -0.0141 | 0.2487 | 0.00001 | -0.5383 | 0 | 0.1058 |
|  | 0.3166 | -0.0654 | 0.2356 | 0.3155 | 0.0017 | 0.2459 | 0.00003 | -0.5911 | 0 | 0.2192 |
| SN(-DC)/SH | 0.3006 | 0.032 | -0.2619 | 0.3462 | 0.0784 | -0.1843 | 0.0005 | 0.44 | 0 | 0.15 |
|  | 0.331 | 0.0516 | -0.2186 | 0.2519 | 0.0487 | -0.3069 | 0.001 | 0.44 | 0 | 0.09 |
|  | 0.3163 | 0.0419 | -0.2413 | 0.3049 | 0.0648 | -0.2507 | 0.00025 | 0.43 | 0 | 0.03 |
| SN(-DC)/FLC | 0.1892 | -0.0243 | -0.3516 | 0.3049 | 0.0648 | -0.2507 | 0.001 | 0.47 | 0 | 0.15 |
|  | 0.0069 | -0.0819 | -0.3915 | 0.2371 | 0.0444 | -0.319 | 0.001 | 0.41 | 0 | 0.44 |
|  | 0.267 | 0.0126 | -0.2976 | 0.3371 | 0.0753 | -0.2018 | 0.001 | 0.46 | 0 | 0.16 |
| SubH/SH | 0.3125 | 0.0395 | -0.2465 | 0.1221 | 0.0104 | -0.3808 | 0.001 | 0.32 | 0 | 0.54 |
|  | 0.3216 | 0.0454 | -0.2335 | 0.0128 | -0.0246 | -0.399 | 0.001 | 0.33 | 0 | 0.8 |
|  | 0.331 | 0.0516 | -0.2186 | 0.1658 | 0.0243 | -0.3632 | 0.001 | 0.35 | 0 | 0.42 |
| SubH/FLC | 0.0628 | -0.0679 | -0.3892 | 0.0312 | -0.0189 | -0.3983 | 0.0005 | 0.38 | 0 | 0.09 |
|  | 0.267 | 0.0126 | -0.2976 | -0.0171 | -0.0336 | -0.3982 | 0.001 | 0.33 | 0 | 0.68 |
|  | -0.1627 | -0.1012 | -0.3511 | 0.1221 | 0.0104 | -0.3808 | 0.0005 | 0.34 | 0 | 0.40 |

---

**Fig S1.**
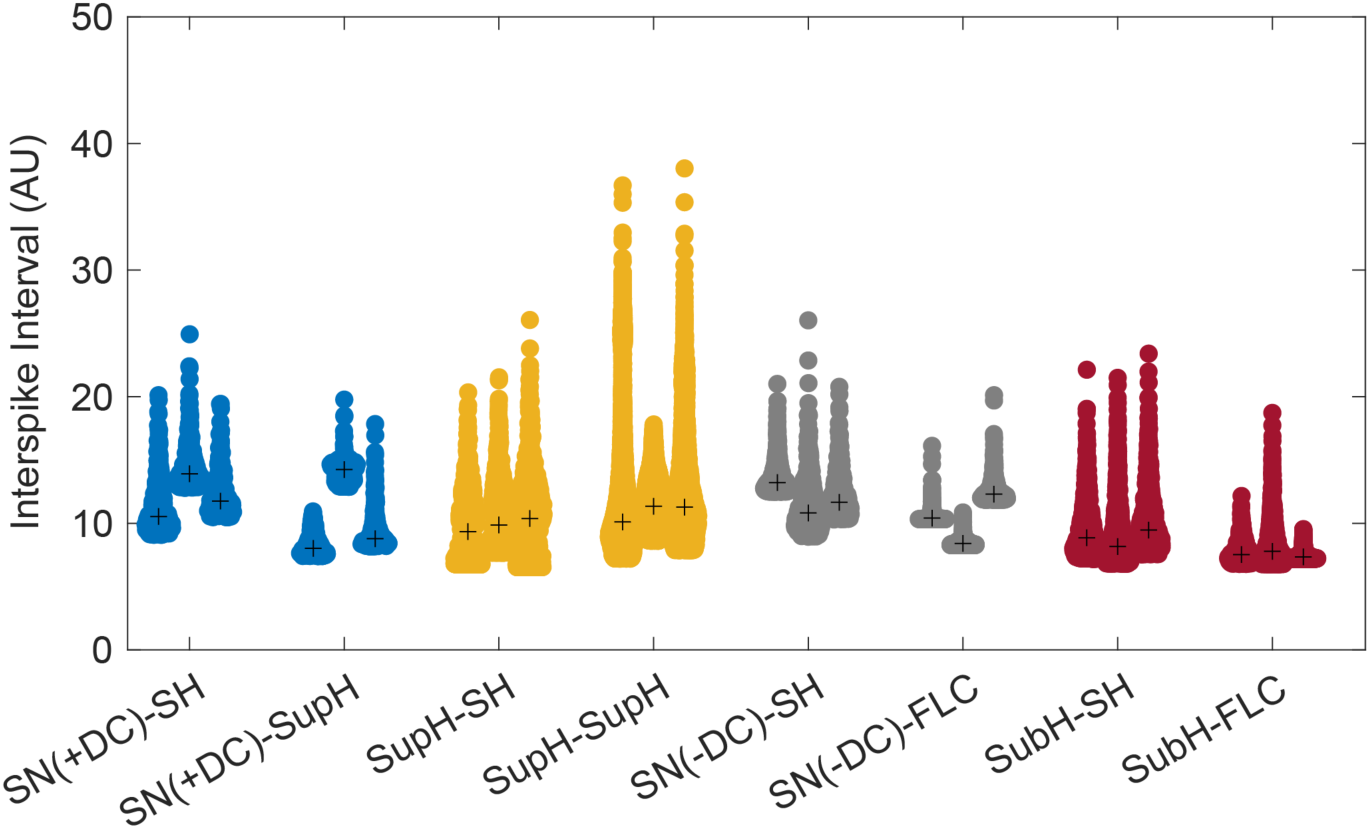
Interspike intervals for each of the 24 node 1 bursting paths implemented. Paths are grouped by dynamotype and colored by onset. Crosses indicate median values. SN(+DC) = saddle node with baseline shift, supH = supercritical Hopf, subH = subcritical Hopf, SN(-DC) = saddle node without baseline shift, SH = saddle homoclinic, FLC = fold limit cycle.

**Fig S2.**
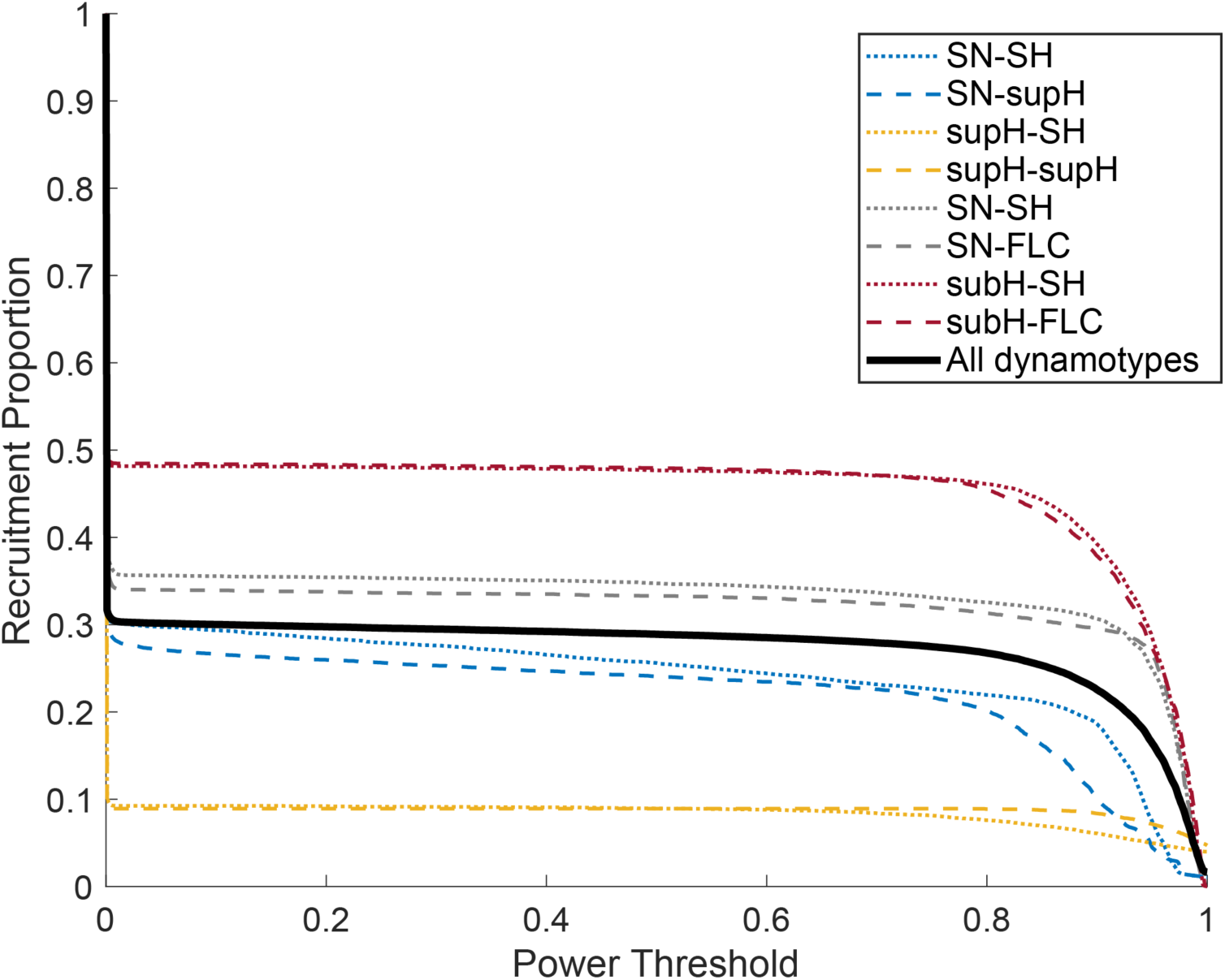
Recruitment proportion as a function of power threshold. Timeseries were high-pass filtered, then mean square power was calculated for each node during the coupling period. Node 2 power was divided by node 1 power and used for identification of recruitment via thresholding. For all dynamotypes, recruitment proportions were consistent for power thresholds between 0.1 and 0.8. Thus, simulations above a power threshold of 0.1 were included in the analyses. SN(+DC) = saddle node with baseline shift, supH = supercritical Hopf, subH = subcritical Hopf, SN(-DC) = saddle node without baseline shift, SH = saddle homoclinic, FLC = fold limit cycle.

**Fig S3.**
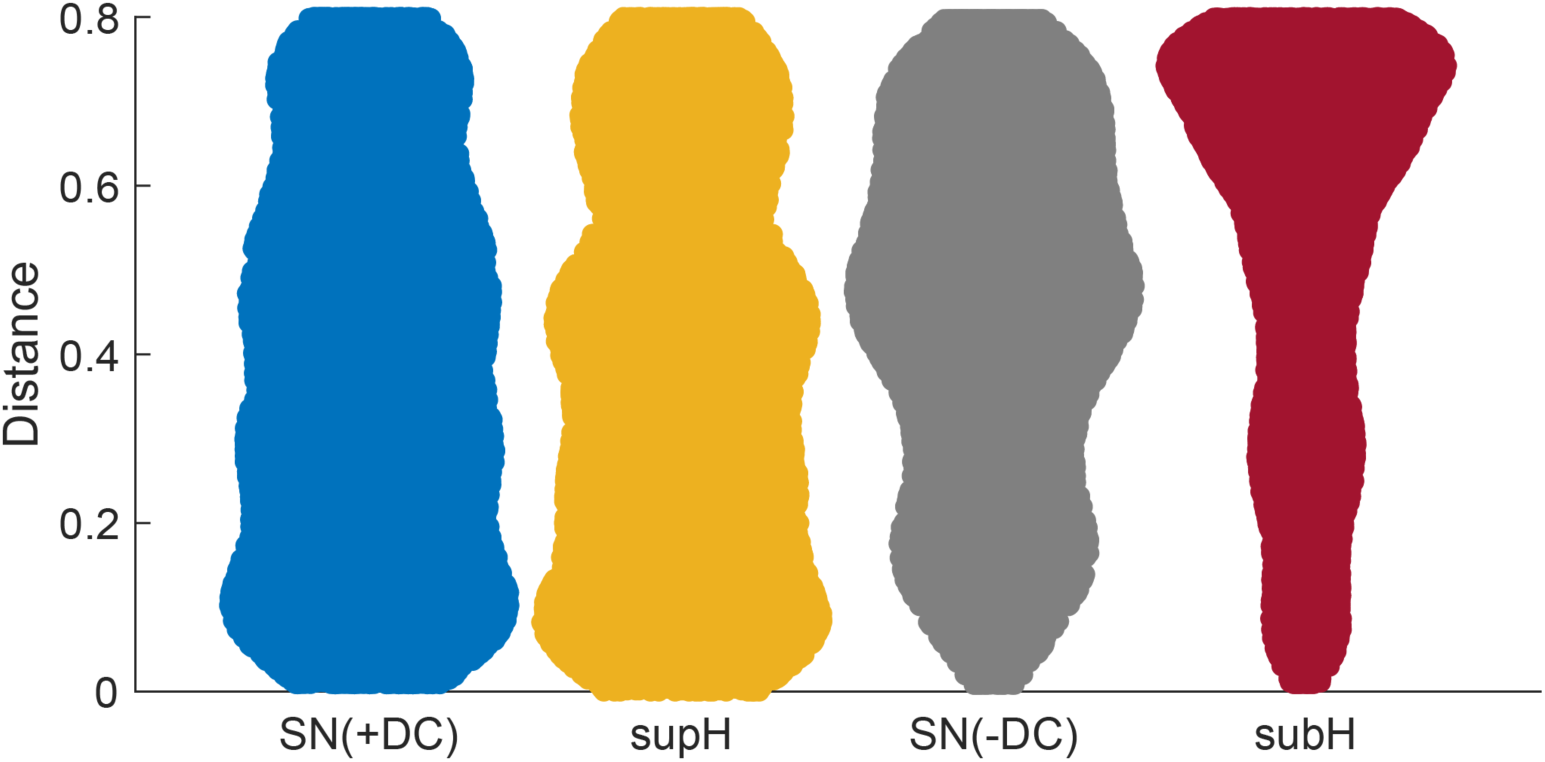
Distributions of distances between node 1 and node 2 at the onset of coupling, with respect to node 1 onset bifurcation. SN(+DC) = saddle node with baseline shift, supH = supercritical Hopf, SN(-DC) = saddle node without baseline shift, subH = subcritical Hopf.

**Fig S4.**
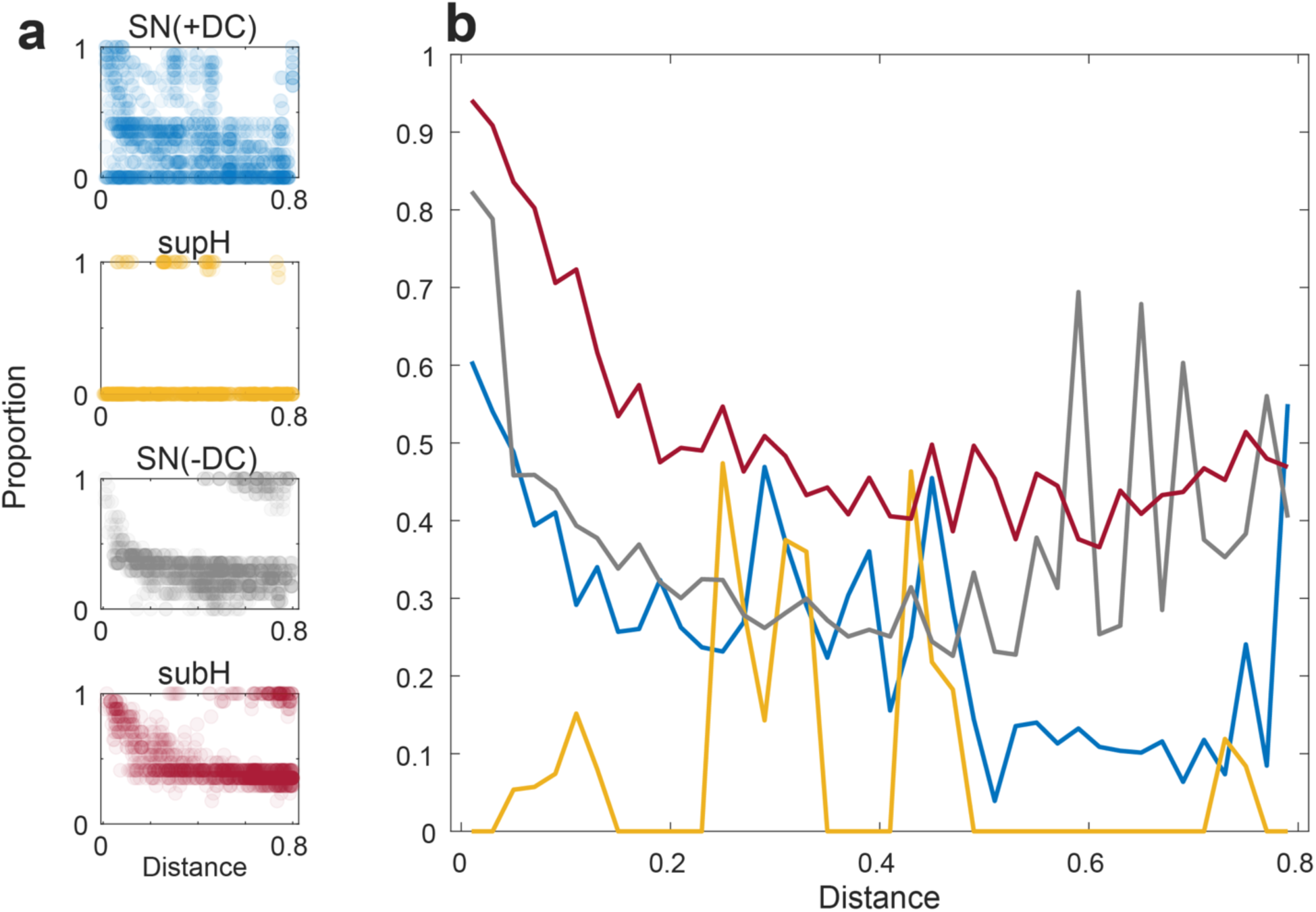
**(a)** Recruitment proportions versus distance between node 1 and node 2 at the onset of coupling with respect to each onset bifurcation. Each point represents one combination of node 1 path and node 2 initial location. SupH-onset dynamotypes recruited a limited number of locations. **(b)** Trends in recruitment proportion. Proportions were computed in 40 equally spaced bins (bin width = 0.02). SN(+DC) = saddle node with baseline shift, supH = supercritical Hopf, SN(-DC) = saddle node without baseline shift, subH = subcritical Hopf.

**Fig S5.**
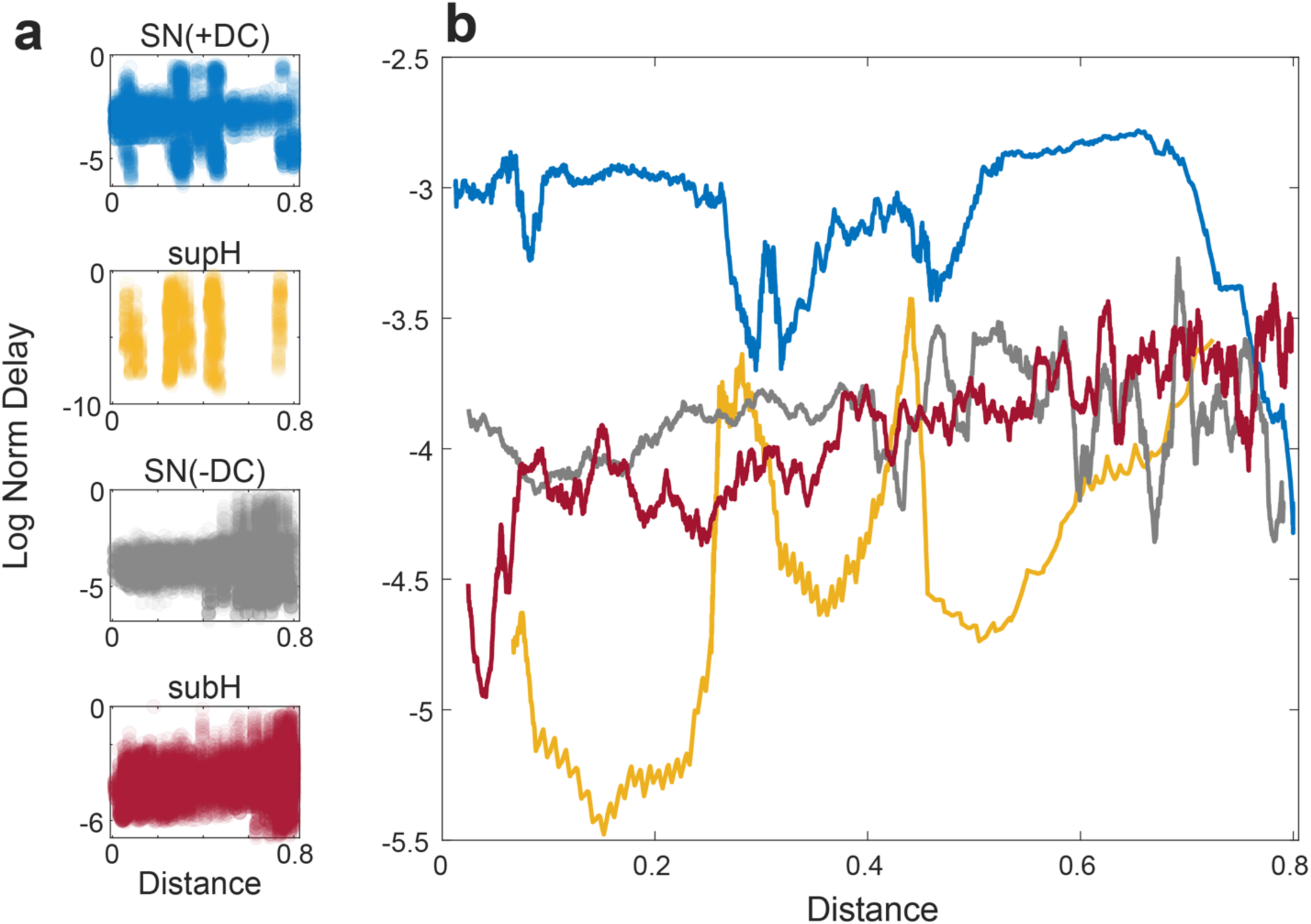
**(a)** Log of normalized recruitment delay versus distance between node 1 and node 2 at the onset of coupling with respect to each onset bifurcation. Each point represents one simulation with recruitment. SupH-onset dynamotypes recruited a limited number of locations. **(b)** Moving averages (window size = 300) of the data in (a). SN(+DC) = saddle node with baseline shift, supH = supercritical Hopf, SN(-DC) = saddle node without baseline shift, subH = subcritical Hopf.

**Fig S6.**
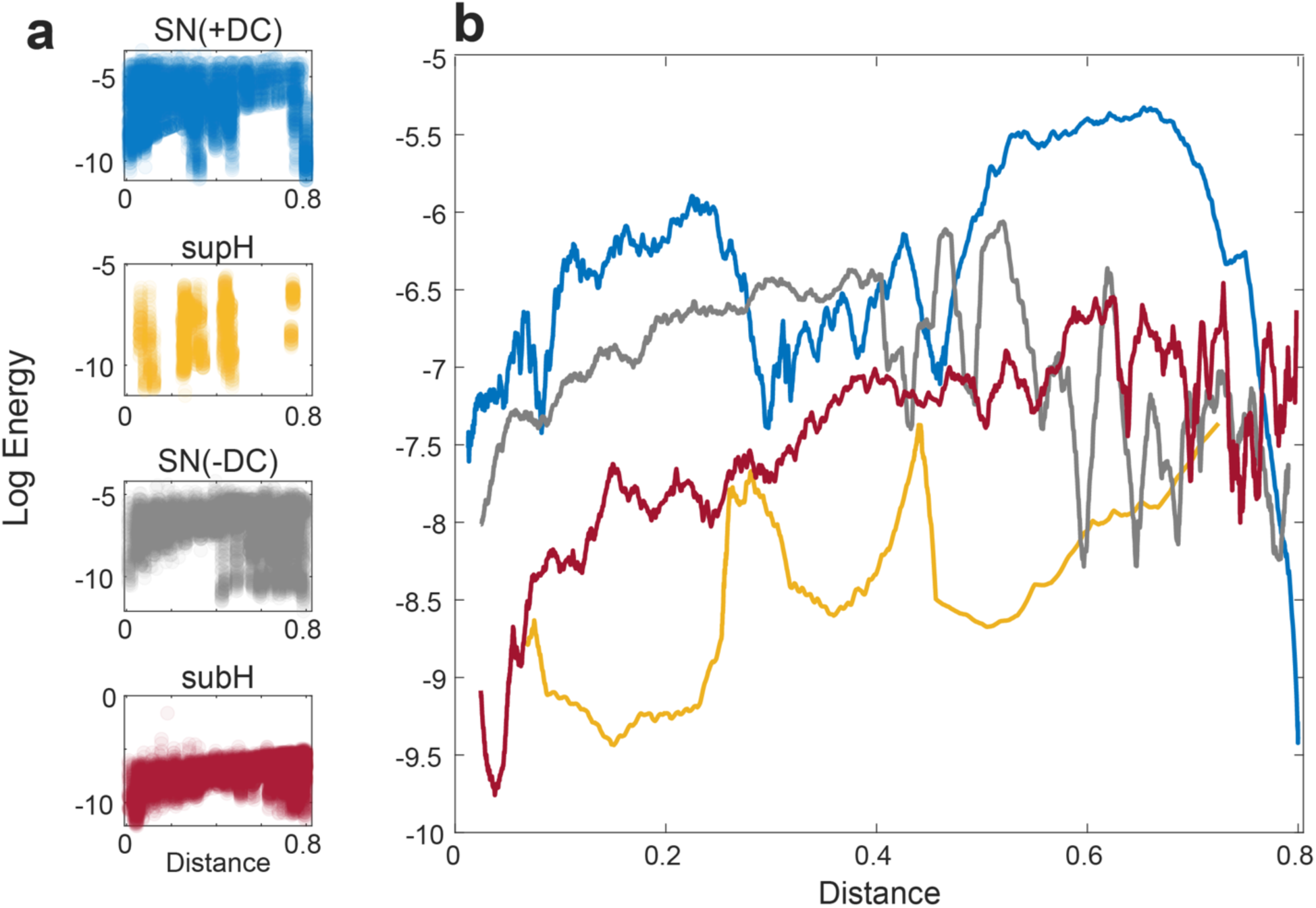
**(a)** Log of recruitment energy versus distance between node 1 and node 2 at the onset of coupling with respect to each onset bifurcation. Each point represents one simulation with recruitment. SupH-onset dynamotypes recruited a limited number of locations. **(b)** Moving averages (window size = 300) of the data in (a). SN(+DC) = saddle node with baseline shift, supH = supercritical Hopf, SN(-DC) = saddle node without baseline shift, subH = subcritical Hopf.

**Fig S7.**
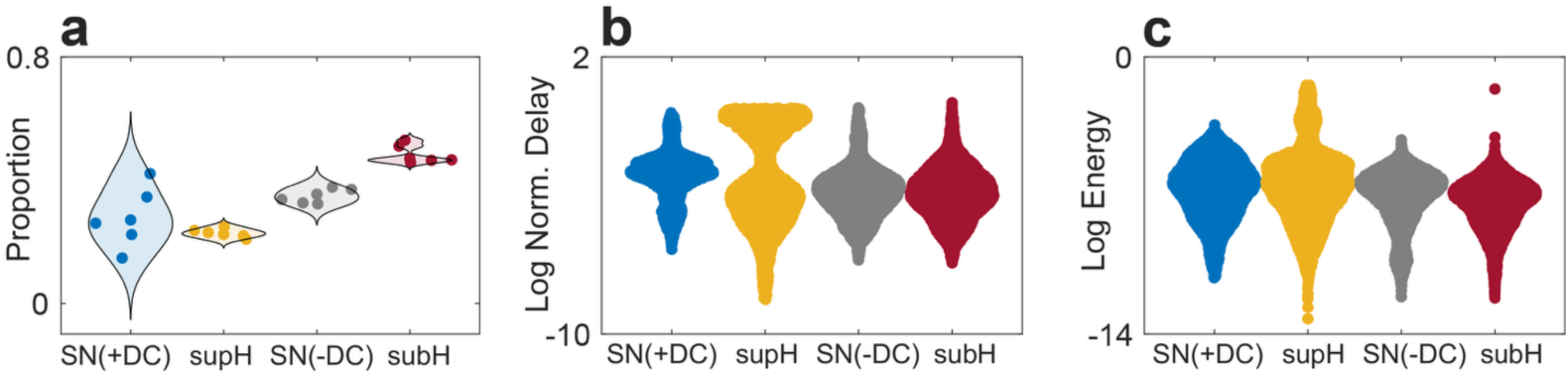
For these results, burst onset was defined as the first departure from rest. Thus, for supH-onset node 1 dynamotypes, coupling began at the preceding SN(+DC) bifurcation. **(a)** Recruitment proportion by node 1 onset. Each datapoint represents one node 1 bursting path. **(b)** Log of normalized recruitment delay and **(c)** log of recruitment energy by node 1 onset. Each datapoint represents one simulation. SN(+DC) = saddle node with baseline shift, supH = supercritical Hopf, SN(-DC) = saddle node without baseline shift, subH = subcritical Hopf.

**Fig S8.**
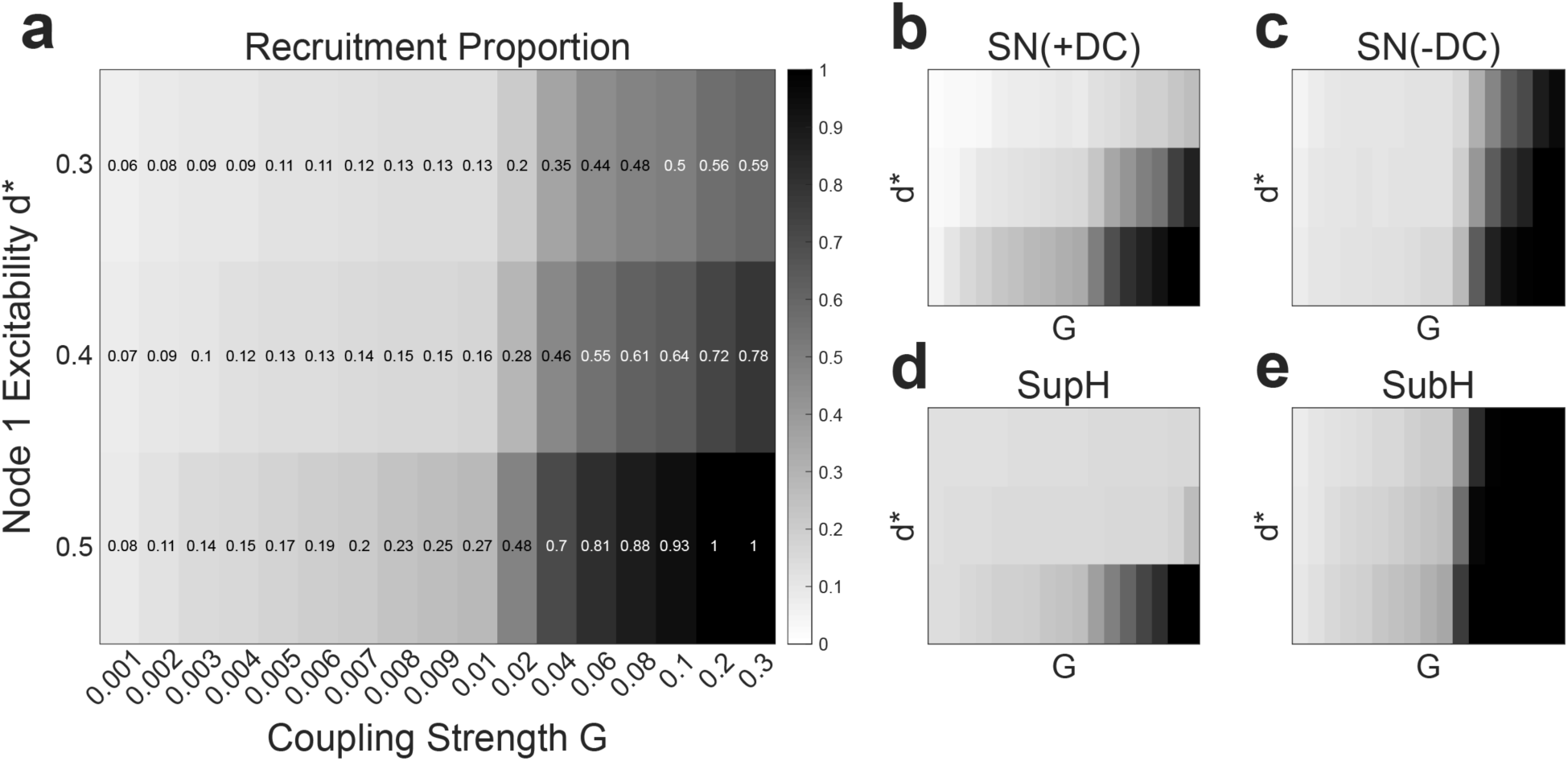
For these results, burst onset was defined as the first departure from rest. Thus, for supH-onset node 1 dynamotypes, coupling began at the preceding SN(+DC) bifurcation. **(a)** Recruitment proportions versus coupling strength (G) and node 1 excitability (d*) for all simulations. Recruitment increased with coupling strength and node 1 excitability. **(b)-(e)** Recruitment proportions versus coupling strength and node 1 excitability for each node 1 onset bifurcation. Dependence on coupling strength and node 1 excitability varied by node 1 onset bifurcation. SN(+DC) = saddle node with baseline shift, supH = supercritical Hopf, SN(-DC) = saddle node without baseline shift, subH = subcritical Hopf.

**Fig S9.**
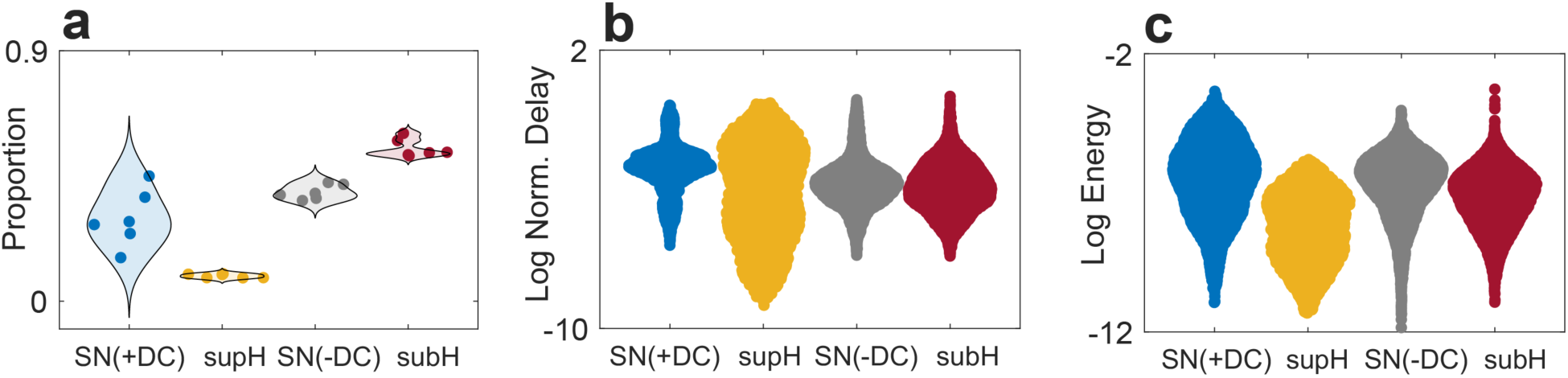
For these results, the coupling term in Eqn. 9 was modified to ensure results were robust to choice of coupling term. **(a)** Recruitment proportion by node 1 onset. Each datapoint represents one node 1 bursting path. **(b)** Log of normalized recruitment delay and **(c)** log of recruitment energy by node 1 onset. Each datapoint represents one simulation. SN(+DC) = saddle node with baseline shift, supH = supercritical Hopf, SN(-DC) = saddle node without baseline shift, subH = subcritical Hopf.

**Fig S10.**
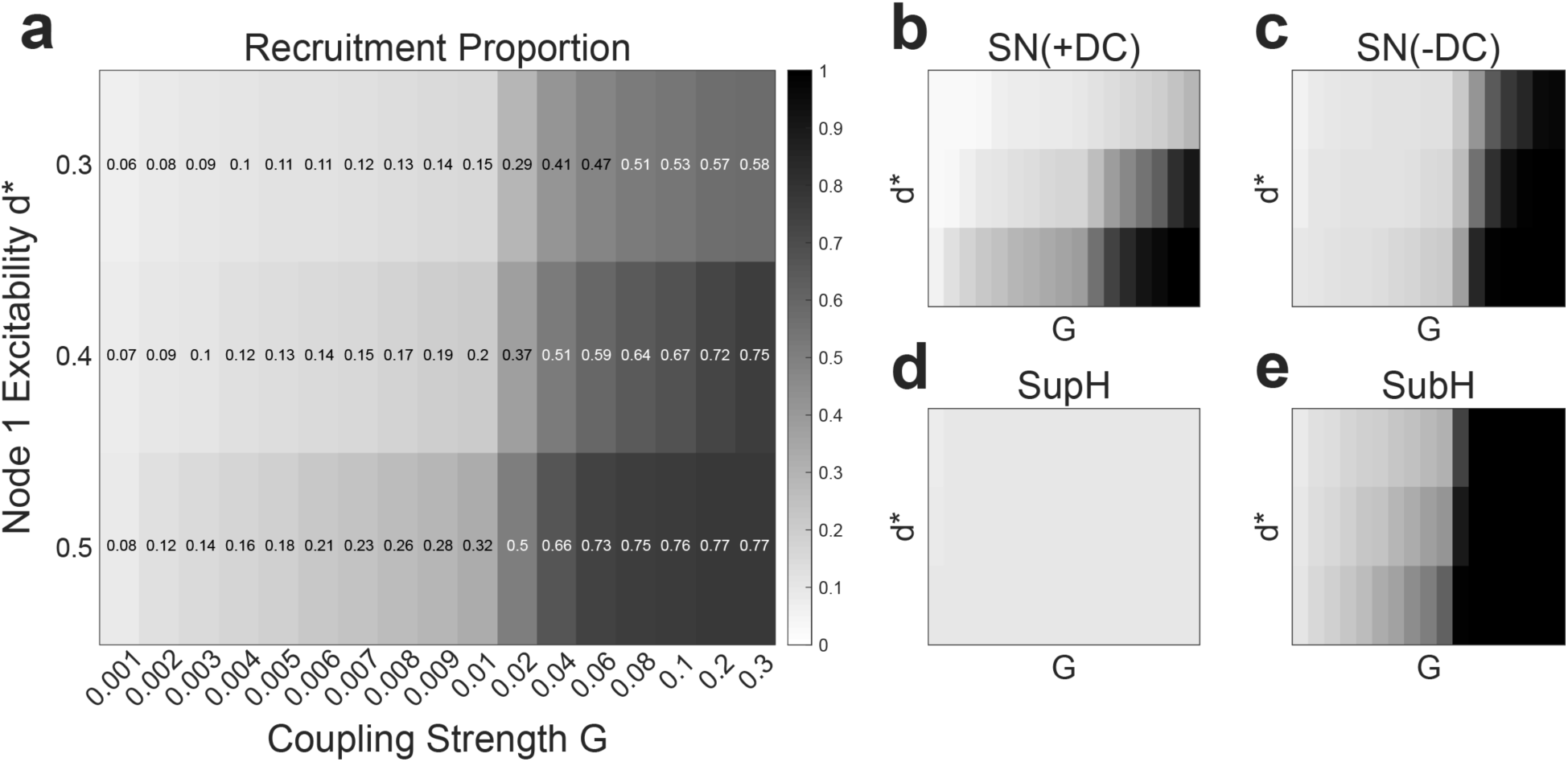
For these results, the coupling term in Eqn. 9 was modified to ensure results were robust to choice of coupling term. **(a)** Recruitment proportions versus coupling strength (G) and node 1 excitability (d*) for all simulations. Recruitment increased with coupling strength and node 1 excitability. **(b)-(e)** Recruitment proportions versus coupling strength and node 1 excitability for each node 1 onset bifurcation. Dependence on coupling strength and node 1 excitability varied by node 1 onset bifurcation. SN(+DC) = saddle node with baseline shift, supH = supercritical Hopf, SN(-DC) = saddle node without baseline shift, subH = subcritical Hopf.

**Fig S11.**
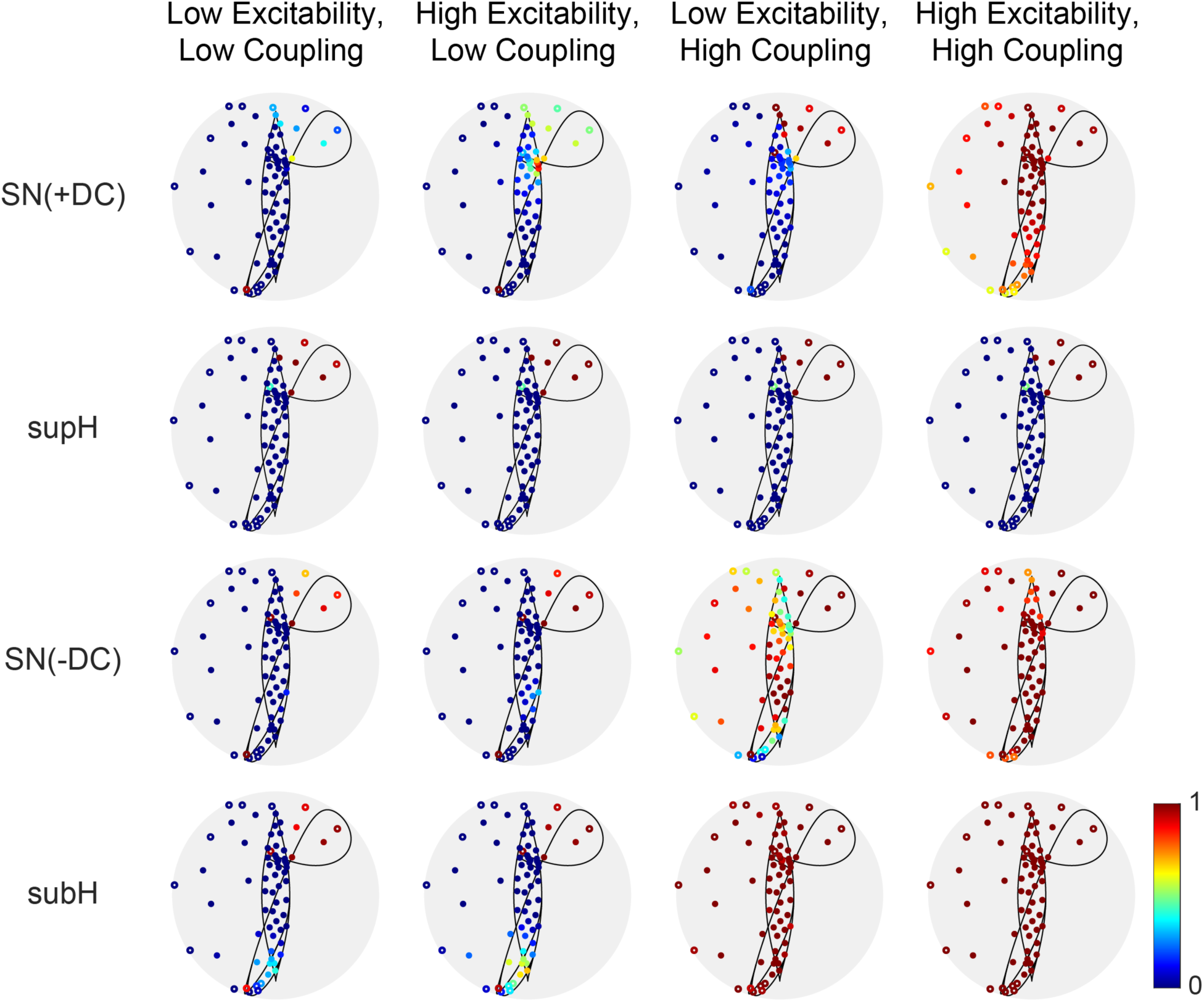
Recruitment proportions for each node 2 initial location, separated by node 1 onset bifurcation and coupling strength and excitability values. Low excitability was d* = 0.3; high excitability was d* = 0.5. Low coupling strength was G < 0.01; high coupling strength was G > 0.05. Low and high coupling strengths were defined based on the observation that overall recruitment proportions increased around a coupling strength of 0.02. SN(+DC) = saddle node with baseline shift; supH = supercritical Hopf; SN(-DC) = saddle node without baseline shift; subH = subcritical Hopf.

**Fig S12.**
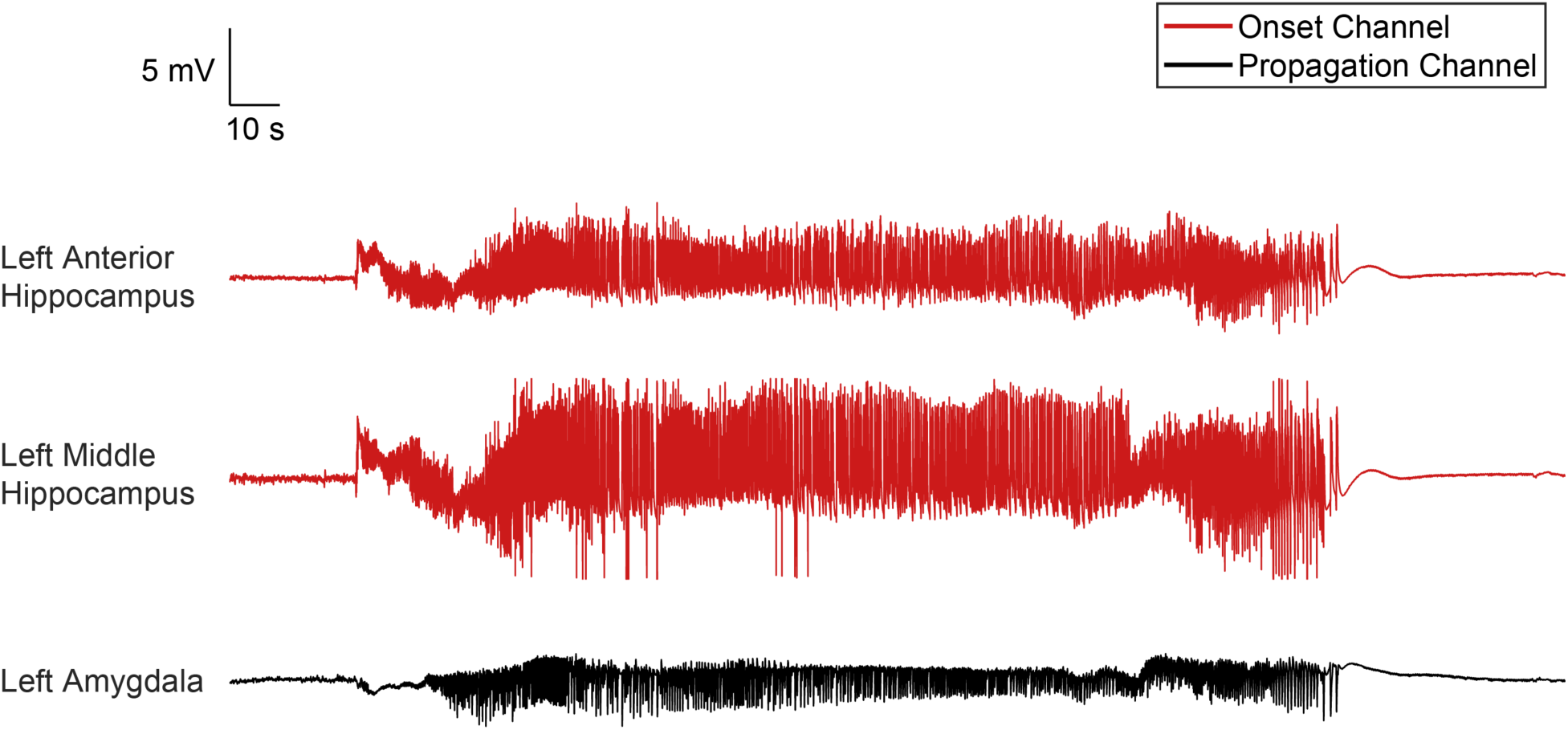
Supplementary seizure propagation example 1. There is a baseline shift at onset in the onset channels, compatible with a SN(+DC) onset bifurcation. In the propagation channel, increasing amplitude at onset is consistent with a supH onset bifurcation.

**Fig S13.**
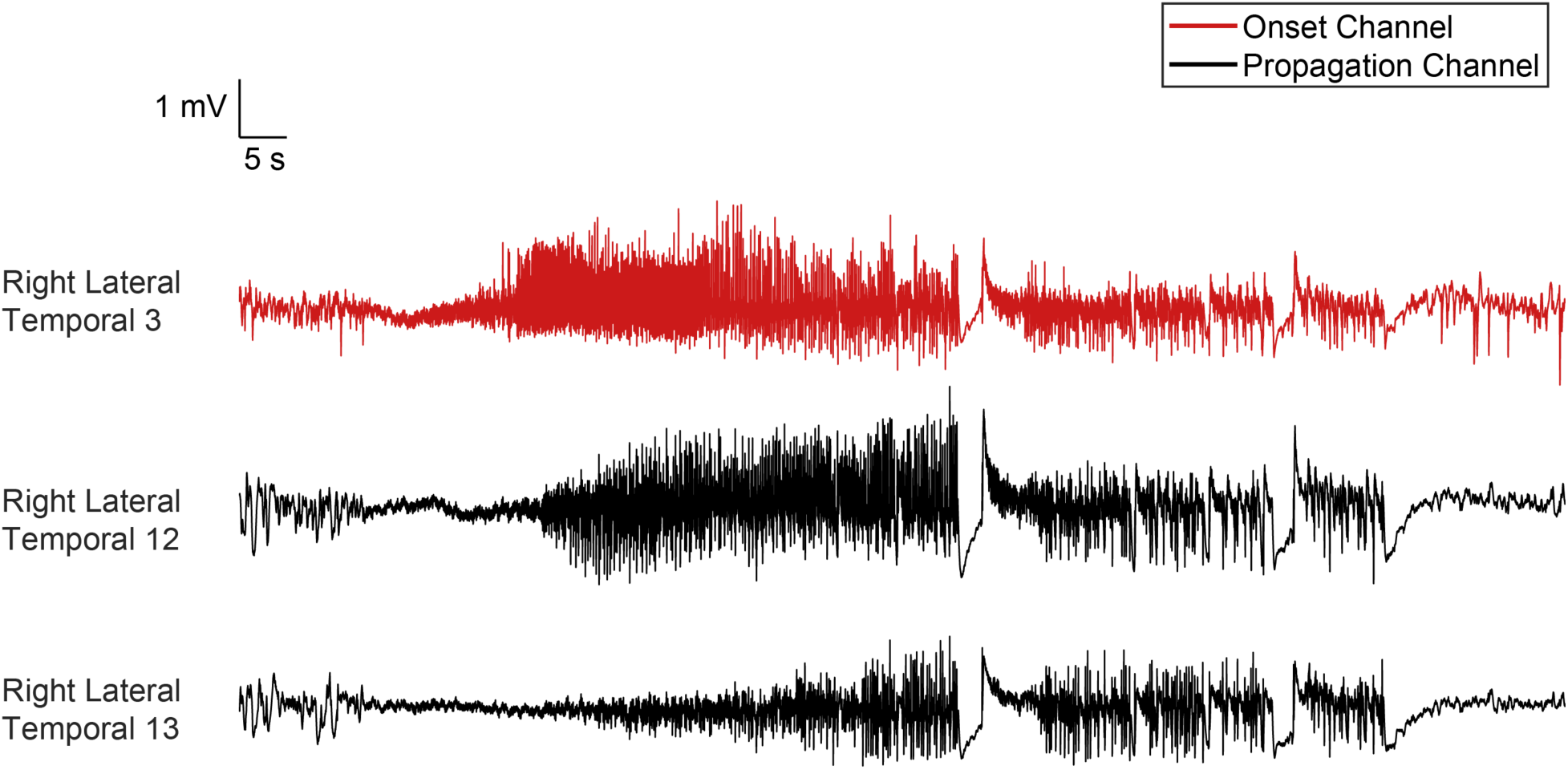
Supplementary seizure propagation example 2. Increasing amplitude at onset suggests supH onset bifurcations in all channels. Numbers indicate electrode contacts; all contacts were located on a grid electrode placed over the right lateral temporal region.

**Fig S14.**
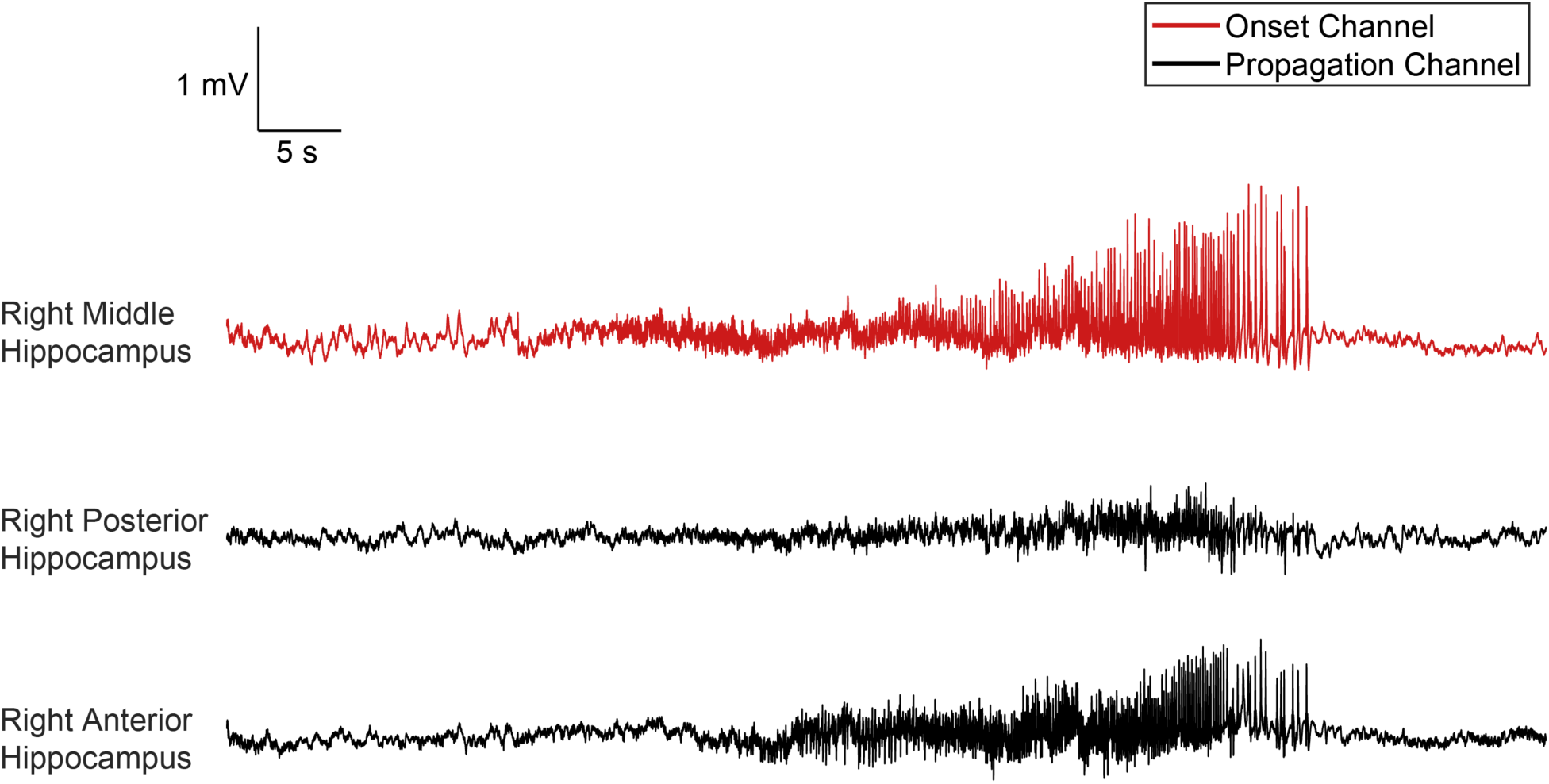
Supplementary seizure propagation example 4. At onset, amplitude is arbitrary in all channels, consistent with subH or SN(-DC) onset bifurcations. Note that although there is increasing amplitude, particularly in the right middle hippocampus, this does not occur until more than 15 seconds after the seizure began. That increasing amplitude was not part of the initial bursting pattern and likely represents volume conduction from neighboring tissue.

